# Bacterial-derived cyclic lipopeptides suppress dauer arrest in *C. elegans*

**DOI:** 10.64898/2026.09.14.751545

**Authors:** Xiao Wang, Kelsie M. Nauta, Marco E. Mechan-Llontop, Darrick R. Gates, Nirmal Chaudhary, Amudha Ramchandran, Kersten K. Sykes, Corrie L. Sakowski, Vashnie H. Hartwell, Jade Tarango, Ashootosh Tripathi, Nicholas O. Burton

## Abstract

Secondary bacterial metabolites can regulate animal physiology, but the specific bacterial genes and molecules responsible for regulating, synthesizing, and transporting these molecules and the mechanisms by which they impact animal physiology often remain unclear. Here, we used *Caenorhabditis elegans* dauer formation as a gross morphological readout to identify bacterial mechanisms that promote development when animal TGF-β or insulin signaling is impaired. We found that the native *C. elegans* microbiome constituent, *Pseudomonas lurida*, suppresses dauer arrest and that this activity requires the two-component system, GacS/GacA, and the downstream *massABC* nonribosomal peptide synthetase cluster. Using sequential chromatography followed by mass spectrometry and NMR, we identified Massetolide F, a bacterial cyclic lipopeptide biosurfactant, as a *massABC*-dependent product. In addition, we found that purified Massetolide F was sufficient to restore dauer suppression to *P. lurida ΔmassBC*. We also found that surfactin C, a cyclic lipopeptide from *B. subtilis*, similarly suppressed dauer formation whereas the cyclic lipopeptide antibiotic daptomycin did not, indicating that dauer suppression is shared by a subset of cyclic lipopeptides rather than being a general property of the class. Lastly, we found that *massABC*-dependent Massetolide F production induces the expression of the acid sphingomyelinase *asm-3* in *C. elegans,* that *asm-3* is required for Massetolide F induced dauer suppression, and that loss of Massetolide F production also leads to reduced bacterial clearance in host animals. Together, these findings identify Massetolide F as a microbiome-derived molecule that promotes animal development and reveal an unexpected function for bacterial biosurfactants in regulating animal physiology.

## Introduction

Microbiome metabolites play diverse roles in regulating animal physiology. For example, in the mammalian gut, microbial short chain fatty acids, bile acid derivatives, tryptophan metabolites, and polyamines regulate intestinal barrier function, immune cell activity, enteroendocrine signaling, glucose metabolism, and inflammatory disease states^1–4^. However, for many host-associated bacteria, the specific bacterial genes, biosynthetic pathways, and metabolites that modify host physiological traits remain unknown^5,6^. A central goal of host microbiome research is therefore to define the molecular mechanism by which host-associated bacteria regulate host physiology and disease^7^.

*Caenorhabditis elegans* provides a powerful system for studying microbiome-metabolite-host interactions, because the animals feed on bacteria, which become both their diet and microbiome, and because the collection of germ-free animals can be performed at scale allowing for high-throughput based approaches that pair an animal model with bacterial monocultures. Previous studies have shown that primary bacterial metabolites alter *C. elegans* development, metabolism, stress resistance, and lifespan^8–13^. For example, vitamin B12 produced by *Comamonas* accelerates development and alters host propionate metabolism, whereas bacterial folate and coenzyme Q metabolism have been linked to changes in aging and lifespan^8–10^. These bacterial-derived molecules are used as nutrients by the host (e.g vitamins), and thus likely alter animal physiology by changing the diet. By contrast, secondary bacterial metabolites, which are not thought to be used as nutrients by the host, have also been reported to impact *C. elegans*. For example, *Pseudomonas* derived cyclic lipopeptides inhibit bacterial pathogens^11,12^. These studies demonstrate that *C. elegans* is a tractable model organism for discovering bacterial mechanisms and metabolites that influence host biology.

Easily assayed biological readouts are needed to use high throughput approaches to identify bacterial species and molecules that impact animal physiology. One example of an easily assayed readout is changes in animal development. *C. elegans* undergo programmed developmental arrests in response to various environmental stressors. Under unfavorable environmental conditions, such as limited food, high population density, and elevated temperature, *C. elegans* larvae enter dauer, a stress resistant and developmentally arrested life stage that can be separated from animals that develop to adulthood^14^. Dauer entry and exit are regulated by conserved TGF-β and insulin signaling pathways^14,15^. In *C. elegans*, TGF-β signaling acts through the DAF-7 ligand, DAF-1/DAF-4 receptors, and downstream SMAD proteins to sense environmental conditions, whereas insulin/IGF-1-like signaling acts through the DAF-2 receptor and a downstream phosphorylation cascade comprising DAF-2, AGE-1/PI3K, PDK-1, AKT kinases, and DAF-16/FOXO to couple nutrient and stress cues to developmental decisions. These pathways are conserved in mammals as TGF-β/SMAD and insulin/IGF-1–PI3K–AKT–FOXO signaling modules, which regulate metabolism, growth, and immune function^15,16^. Loss of function mutations in these pathways cause dauer constitutive phenotypes at elevated temperatures. We hypothesized that unidentified bioactive bacterial metabolites lead to increased animal TGF-β and insulin signaling or alleviate the consequences of reduced TGF-β and insulin signaling and thus alter the developmental trajectory of these mutant animals. The overall goal of this work is to identify novel small molecules that regulate TGF-β and insulin signaling in animals that then may be optimized to modulate metabolic, fibrotic, inflammatory, or cancer associated disease mechanisms which have been previously linked to these signaling pathways.

We report that *Pseudomonas lurida* suppresses dauer arrest in multiple *C. elegans* TGF-β and insulin signaling mutant models. We further report that *P. lurida* uses a regulatory axis in which the GacS/GacA two-component system activates the downstream *massABC* nonribosomal peptide synthetase cluster, as previously reported in the closely related *Pseudomonad*, *P. flourescens*^17^. Activation of the *massABC* biosynthetic gene cluster results in the production of Massetolide F which activates the expression of the sphingomyelinase *asm-3* in *C. elegans.* We found that *asm-3* expression is required to suppress *daf-8* dauer formation. Massetolides are cyclic lipopeptide biosurfactants produced by *Pseudomonas* species. Cyclic lipopeptides are composed of a lipid tail linked to a cyclic oligopeptide and are synthesized by a diverse organism including *Aspergillus*, *Streptomyces*, *Pseudomonas*, and *Bacillus*^18–21^. Here, we report that Massetolide F suppression of dauer arrest is not specific to Massetolide F but is also observed for surfactin, a structurally distinct cyclic lipopeptide from *Bacillus subtilis.* Bacterial cyclic lipopeptides have been studied mainly for their microbial and ecological functions, including surface motility, biofilm formation, interspecies competition, plant associated biocontrol, and antimicrobial activity^18,22,23^. These activities have led biosurfactants to be viewed largely as molecules that shape microbial surface behavior, interactions between microbes, and colonization of the host. Our results suggest that in addition to these functions, cyclic lipopeptides such as Massetolide F and surfactin also play a key role in modulating animal insulin and TGF-β signaling. Furthermore, because *P. lurida* was previously found to colonize wild *C. elegans* intestinal microbiomes, our data suggest that this mechanism might play an important role in regulating this previously established host-microbiome relationship^11,24–28^.

## Results

### *Pseudomonas lurida* suppresses dauer arrest in TGF-β and insulin signaling mutants

To identify bacterial isolates that improve developmental defects caused by impaired TGF-β signaling, we screened a library of approximately 4,000 wild bacterial isolates using *daf-1(m40)* animals (**Fig. 1A**), which normally arrest as dauers at 26 °C (**Fig. 1B**), similarly to previous work^13^. Using this approach, we isolated and sequenced a strain of *Pseudomonas lurida* that we designated as NOBb261. This strain promoted a fraction of *daf-1(m40)* animals to develop to adulthood at 26 °C rather than remaining arrested in dauer (**Fig. 1B**). We next examined whether dauer suppression was specific to *daf-1(m40)* or whether *P. lurida* NOBb261 could also suppress dauer arrest in other dauer constitutive (*daf-c*) mutants. Across a panel of mutants in the TGF-β and insulin signaling pathways, *P. lurida* NOBb261 produced mild (<10% adult animals) to moderate (10–30% adult animals) suppression of dauer arrest (**Fig. 1C**) under conditions where >99% of mutant animals normally develop as dauers. The ability of *P*.

**Figure 1.**
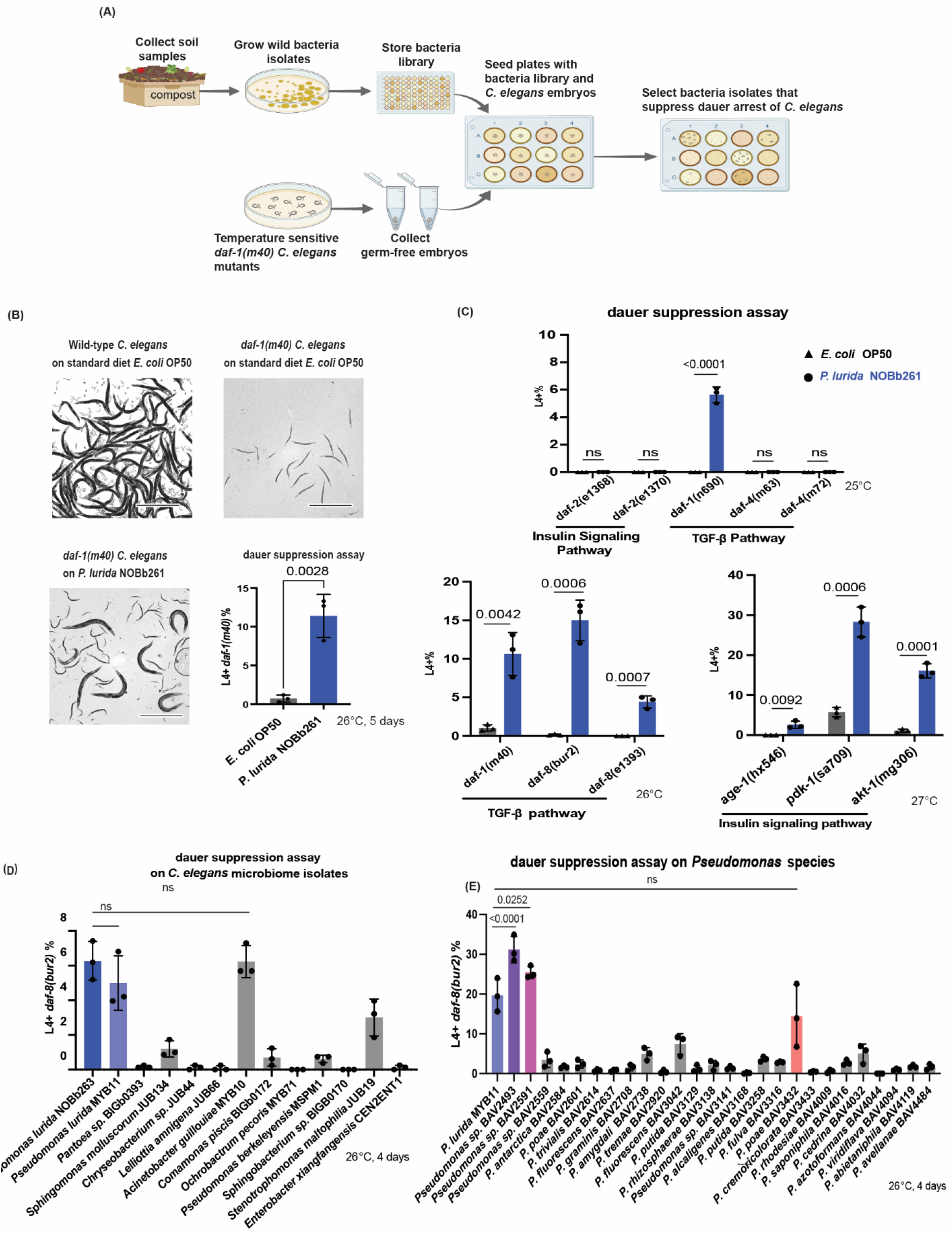
*Pseudomonas lurida* suppresses dauer arrest in TGF-β and insulin signaling mutants. **(A)** Graphical representation of the platform used to screen wild bacterial isolates for suppression of dauer arrest in *daf-1(m40)* mutant *C. elegans*. Germ-free embryos were arrayed on a library of 4,000 wild bacterial isolates and screened for development to the L4 stage or later. Candidate bacterial isolates were retested on 90 mm plates with approximately 500 embryos. Image created in BioRender. **(B)** *Pseudomonas lurida* NOBb261 promotes development of *daf-1(m40)* animals from dauer arrest. Representative images show wild-type *C. elegans* on *E. coli* OP50, and *daf-1(m40)* mutant *C. elegans* on *E. coli* OP50 or *P. lurida* NOBb261. Animals were incubated at 26 °C for 5 days on their respective diets, and the percentage of *daf-1(m40)* animals that developed to the L4 stage or later was quantified. Experiments were performed in triplicate with approximately 500 animals per replicate. An unpaired t-test was used to determine statistical significance. Error bars indicate s.d. from the mean. Exact p values are shown on the bar graph. Scale bar = 1 mm. **(C)** *P. lurida* NOBb261 promotes development of *daf-1(n690)*, *daf-1(m40)*, *daf-8(bur2)*, *daf-8(e1393)*, *akt-1(mg306)*, *age-1(hx546)*, and *pdk-1(sa709)* animals from dauer arrest, but does not promote development of *daf-2(e1368)*, *daf-2(e1370)*, *daf-4(m63)*, or *daf-4(m72)* animals. At 25 °C, the insulin signaling mutants *daf-2(e1368)* and *daf-2(e1370)* and the TGF-β signaling mutants *daf-1(n690)*, *daf-4(m63)*, and *daf-4(m72)* were incubated for 5 days. At 26 °C, the TGF-β signaling mutants *daf-8(bur2)*, *daf-8(e1393)*, and *daf-1(m40)* were incubated for 4, 5, and 5 days, respectively. At 27 °C, the insulin signaling mutants *akt-1(mg306)*, *age-1(hx546)*, and *pdk-1(sa709)* were incubated for 3, 5, and 5 days, respectively. Incubation time and temperature were optimized to induce robust dauer arrest on the *E. coli* OP50 control diet. Experiments were performed in triplicate with approximately 500 animals per replicate. An unpaired t-test was used to determine statistical significance for each dauer constitutive (*daf-c*) strain. Error bars indicate s.d. from the mean. Exact p values are shown on the bar graph; ns, not significant. **(D)** *P. lurida* MYB11 promotes *daf-8(bur2)* development to a similar extent as *P. lurida* NOBb261. Among the *C. elegans* microbiome strains screened from the CGC, *P. lurida* MYB11 and *Acinetobacter guillouiae* MYB10 suppressed *daf-8(bur2)* dauer arrest to a similar extent as *P. lurida* NOBb261. *daf-8(bur2)* animals were incubated at 26 °C for 4 days on their respective diets, and the percentage of animals that developed to the L4 stage or later was quantified. An ordinary one-way ANOVA followed by Sidak’s multiple comparisons test was used to determine statistical significance. Error bars indicate s.d. from the mean; ns, not significant. **(E)** Promotion of development from dauer arrest is not a common feature of the *Pseudomonas* genus. Among 26 non-*lurida Pseudomonas* isolates tested, *Pseudomonas* sp. BAV2493 and *Pseudomonas* sp. BAV2591 promoted higher development from dauer arrest in *daf-8(bur2)* animals, while *P. poae* BAV3432 promoted development at a level similar to *P. lurida* MYB11. Experimental setup and statistical analysis were the same as in panel D.

*lurida* NOBb261 to promote development was not observed in all *daf-c* backgrounds. In particular, strong dauer constitutive alleles (∼100% dauer arrest at 25 °C) in the insulin signaling pathway, including *daf-2(e1368)* and *daf-2(e1370)*, and in the TGF-β pathway, including *daf- 4(m63)* and *daf-4(m72)*, remained arrested as dauers on *P. lurida* NOBb261. These results suggest that *P. lurida* NOBb261 can partially compensate for reduced dauer regulatory signaling but cannot overcome more severe genetic disruptions of these pathways that result in 100% dauer development at lower temperatures (25 °C). Among these strains, *daf-8(bur2)* showed the most consistent response to *P. lurida* NOBb261, with low background development on the normal diet, *E. coli* OP50. The *daf-8(bur2)* allele was identified in the Burton laboratory as a background mutation in an N2-derived strain with an unexpected *daf-c* phenotype. This allele contains a missense mutation that results in threonine to proline substitution at residue 359 of daf-8 (T359P), within the MH2 domain (aa 349–546; UniProtKB Q21733) of the DAF-8 R-SMAD^29^. For this reason, we used *daf-8(bur2)* for subsequent dauer suppression assays.

*Pseudomonas lurida* was previously found to colonize *C. elegans* intestinal microbiome in wild *C. elegans* and an isolate of *P. lurida* (MYb11) is included in the CeMBio *C. elegans* microbiome collection^24,27^. This finding suggests that the effects of *P. lurida* on dauer development might be relevant to its natural ecological interactions with *C. elegans*. We thus tested whether any of the CeMBio isolates of bacteria naturally associated with *C. elegans* microbiomes, including the

MYb11 isolate, also caused suppression of dauer in our models. Consistent with our previous findings, we found that the *P. lurida* strain MYb11 suppressed *daf-8(bur2)* dauer arrest to a similar extent as *P. lurida* NOBb261 (**Fig. 1D**). In addition, we found that while some other CeMBio isolates also mildly suppressed dauer development (*Acinetobacter guillouiae* MYb10 and *Stenotrophomonas sp*. JUb19), most CeMBio bacteria did not suppress dauer development when compared to *E. coli* OP50. We also asked whether dauer suppression was a general feature of the *Pseudomonas* genus or if it was unique to specific species or isolates. To address this, we tested 26 additional environmental *Pseudomonas* species^30^ for their ability to suppress dauer arrest in *daf-8(bur2)* animals at 26 °C. Only 3 of the 26 non-*lurida Pseudomonas* isolates tested, *Pseudomonas* sp. BAV2493, *Pseudomonas* sp. BAV2591, and *P. poae* BAV3432, suppressed dauer arrest at levels similar to or greater than *P. lurida* NOBb261 (**Fig. 1E**). Thus, dauer suppression is not a common feature of the *Pseudomonas* genus or to all CeMBio collection isolates naturally associated with the *C. elegans* microbiome. Because MYb11 is publicly available and has been previously sequenced and studied, we used this strain for subsequent dauer suppression assays and genetic manipulations.

### GacA regulated massetolide production is required in *P. lurida* to suppress dauer arrest

To identify bacterial mechanisms that allow *P. lurida* to suppress dauer arrest in *daf-8(bur2)* animals, we performed a transposon mutagenesis screen to identify bacterial genes and pathways required for dauer arrest suppression (**Fig. 2A**). From 3000 screened random transposon mutants of *P. lurida*, we identified two strains that had mutations in the same gene, *gacA. gacA* encodes part of the GacS/GacA two component system^31,32^. Both mutants failed to suppress dauer arrest in *daf-8(bur2)* animals. Based on these results, we hypothesized that *gacA* is required in *P. lurida* to suppress dauer arrest. To test this hypothesis, we generated a clean, in-frame knock out of *gacA* in *P. lurida* MYb11 and complemented the deletion mutant with a wild type copy of *gacA*. We then tested if the deletion mutants phenocopied the transposon mutants on dauer arrest in *daf-8(bur2)* animals. Deletion of *gacA* abolished dauer suppression, as *daf-8(bur2)* animals fed *P. lurida* MYb11 *ΔgacA* showed almost no development to adulthood. By contrast, complementation with WT *gacA* restored dauer suppression, with 10 to 15% of animals developing to adulthood, similar to animals fed wild type *P. lurida* MYb11 (**Fig. 2B**). We conclude that *gacA* is required in *P. lurida* MYb11 to suppress dauer arrest in *daf-8(bur2)* animals.

**Figure 2.**
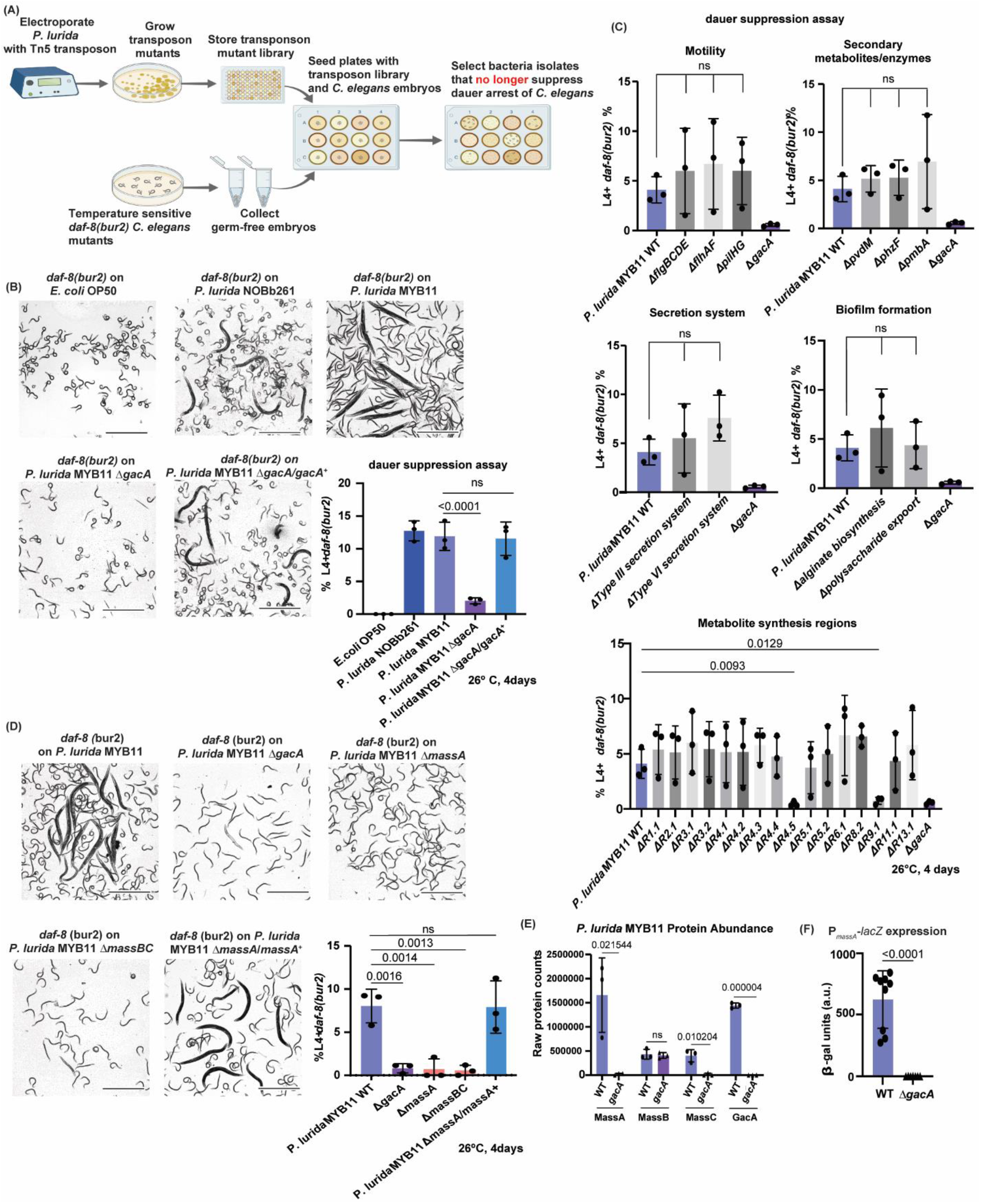
GacA-regulated Massetolide biosynthesis is required for *P. lurida*-mediated dauer suppression. **(A)** Graphical representation of the transposon mutagenesis screen used to identify *P. lurida* genes required for dauer suppression. *P. lurida* was mutagenized by Tn5 transposon insertion, and individual transposon mutants were grown, archived, and screened on plates seeded with germ-free *daf-8(bur2)* embryos. Candidate bacterial mutants were selected based on loss of the ability to promote *C. elegans* development. Image created in BioRender. **(B)** *gacA* is required for *P. lurida* MYB11-mediated suppression of *daf-8(bur2)* dauer arrest. Representative images show *daf-8(bur2)* animals grown on *E. coli* OP50, *P. lurida* NOBb261, *P. lurida* MYB11, *P. lurida* MYB11 Δ*gacA*, or *P. lurida* MYB11 Δ*gacA* complemented with wild-type *gacA*. Animals were incubated at 26 °C for 4 days, and the percentage of animals that developed to the L4 stage or later was quantified(see graph). Experiments were performed in triplicate with approximately 500 animals per replicate. Statistical significance was determined by ordinary one-way ANOVA followed by Sidak’s multiple comparisons testing. Error bars indicate s.d. from the mean. Exact p values are shown on the graph; ns, not significant. Scale bar = 1 mm. **(C)** Screening of targeted *P. lurida* MYB11 deletion mutants for loss of dauer suppression. Mutants affecting motility, secondary metabolite or enzyme-associated genes, secretion systems, biofilm-associated pathways, and predicted metabolite synthesis regions were tested for their ability to suppress dauer arrest in *daf-8(bur2)* animals at 26 °C for 4 days (Supplemental Table 4). Deletion of *gacA* was included as a negative control. Experiments were performed in triplicate with approximately 500 animals per replicate. Statistical significance was determined by ordinary one-way ANOVA followed by Sidak’s multiple comparisons testing within each group. Error bars indicate s.d. from the mean. Exact p values are shown on the graphs; ns, not significant. Each graph presents the same WT data and the mutants were tested in the same experiments. **(D)** The *massABC* biosynthetic cluster (correlated to ΔR4.5, ΔR9.1) is required for *P. lurida* MYB11-mediated dauer suppression. Representative images show *daf-8(bur2)* animals grown on WT *P. lurida* MYB11, *P. lurida* MYB11 Δ*gacA*, *P. lurida* MYB11 Δ*massA*, *P. lurida* MYB11 Δ*massBC*, or *P. lurida* MYB11 Δ*massA* complemented with WT *massA*. Animals were incubated at 26 °C for 4 days, and the percentage of animals that developed to the L4 stage or later was quantified. Experiments were performed in triplicate with approximately 500 animals per replicate. Statistical significance was determined by ordinary one-way ANOVA followed by Sidak’s multiple comparisons testing. Error bars indicate s.d. from the mean. Exact p values are shown on the graph; ns, not significant. **(E)** Proteomic analysis of WT *P. lurida* MYB11 and *P. lurida* MYB11 Δ*gacA*. Raw protein counts are shown for MassABC, and GacA. Statistical significance was determined using an unpaired t-test. Error bars indicate s.d. from the mean. Exact p values are shown on the graph; ns, not significant. **(F)** *gacA* is required for activation of a *massA* reporter in *P. lurida* MYB11. β-galactosidase activity was measured in wild-type *P. lurida* MYB11 and *P. lurida* MYB11 Δ*gacA* carrying the P*_massA_*-lacZ reporter. Loss of *gacA* strongly reduced reporter activity (reflected as β-gal units). Statistical significance was determined using an unpaired t-test. Error bars indicate s.d. from the mean. Exact p value is shown on the graph.

The GacS/GacA two component system has been reported to function as a post transcriptional regulator of multiple downstream pathways, including secretion systems, biofilm formation, and secondary metabolite production, depending on the bacterial species^17,31–33^. We therefore hypothesized that a specific downstream pathway or metabolite regulated by GacS/GacA is required for *P. lurida* to suppress dauer arrest. We first deleted genes associated with several established downstream functions regulated by the GacS/GacA system in other *Pseudomonas sp.* including genes required for bacterial motility, biofilm formation, and various secretion systems. We found that none of these functions were required for *P. lurida* to suppress dauer (Fig. 2C). To identify candidate downstream pathways that might be regulated by GacS/GacA, we used antiSMASH^34^ to predict operons in *P. lurida* that would encode secondary metabolite biosynthetic clusters which are also potentially regulated by GacS/GacA (**Table 1**). We then tested these *P. lurida* MYb11 mutants for their ability to suppress dauer arrest in *daf-8(bur2)* animals at 26 °C (**Fig. 2C**). Among the mutants tested, we found that a cluster of nonribosomal peptide synthetases predicted to produce the cyclic lipopeptide massetolide^35^ was required for dauer suppression (**Supplemental Fig. 1**). Deletion of these genes, *massA* or *massBC,* resulted in almost complete loss of dauer suppression, whereas complementation of the *massA* deletion mutant with a wild type copy of *massA* restored dauer suppression (**Fig. 2D**). These results suggest that the *massABC* biosynthetic cluster is required for *P. lurida* MYB11 to suppress dauer arrest.

**Table 1:**
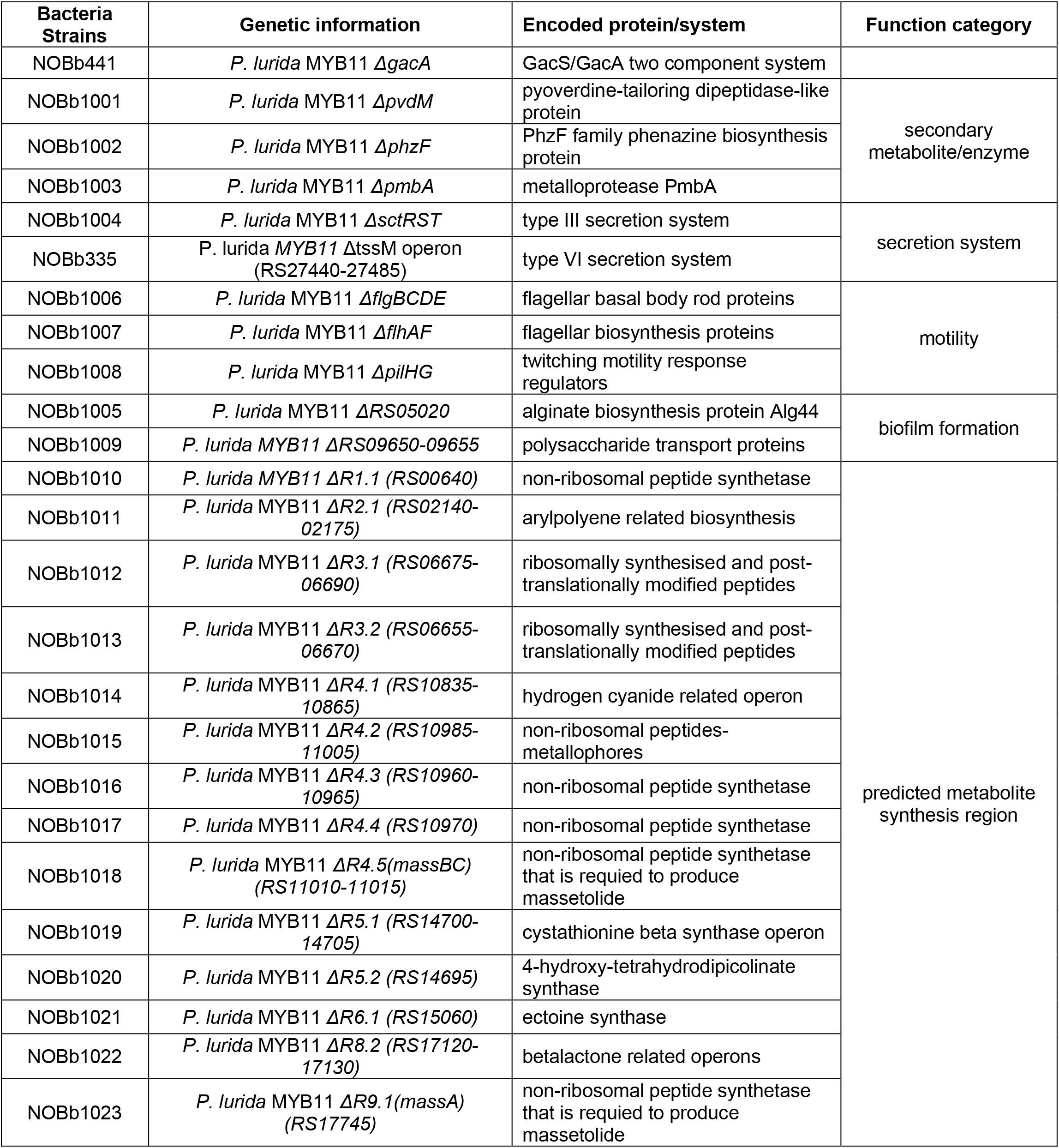
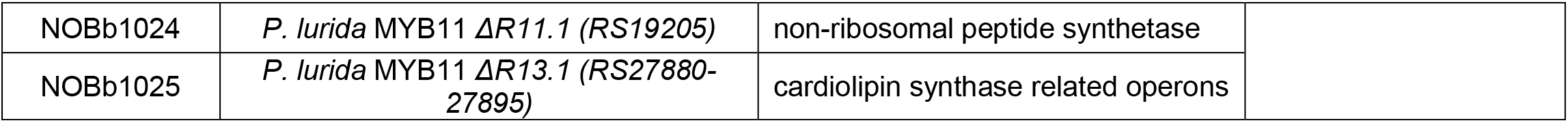
Bacteria knockouts tested for effects on dauer suppression.

To confirm whether *massABC* is regulated by the GacS/GacA two component system, we compared the MassABC protein abundance in WT *P. lurida* MYb11 to MYb11 *ΔgacA* (**Fig. 2E and Supplementary Fig. 2**). This analysis revealed significantly less MassA and MassB protein abundance in MYb11 Δ*gacA* than in WT (**Fig. 2E**). Since *gacA* is known to regulate the small RNAs *rsmY* and *rsmZ*, which function by binding to and inhibiting mRNA^17,31,36^, and to validate our global proteomics findings, we built a translational reporter by fusing the native promoter and first 180 bps of *massA* to a promoterless *lacZ* (*P_massA_’-‘lacZ*). We found activation of P*_massA_’- ‘lacZ was dependent on gacA* (**Fig. 2F**).Together, these findings establish a regulatory axis in *P. lurida* in which the upstream GacS/GacA two component system activates the downstream *massABC* nonribosomal peptide synthetase cluster. This pathway is required for production of massetolide class cyclic lipopeptide(s) and is essential for *P. lurida* MYb11 to suppress dauer arrest and promote development in *daf-8(bur2)* animals.

### GacA-regulated massetolide production is required for *P. lurida* motility and intestinal clearance by the host

Massetolides are a class of cyclic lipopeptide biosurfactants reported to contribute to bacterial surface motility, biofilm formation, and microbial competition^16–18^. We therefore hypothesized that disruption of the *gacS/gacA**–**massABC* regulatory axis and the resulting loss of massetolide production would affect *P. lurida* motility and intestinal clearance by the host. To test this hypothesis, we assayed *P. lurida* motility mediated by the flagella (swarming and swimming) and by the pili (twitching) (**Supplemental Fig. 3**). We found *ΔmassA* and *ΔmassBC* severely impaired both flagella-mediated motilities (**Supplemental Fig. 3**) and, surprisingly, pili-mediated motility (**Supplemental Fig. 3**). Notably, *P. lurida ΔflgBCDE* did not disrupt dauer suppression (**Supplemental Fig. 3**), so we conclude that loss of flagella-mediated motility because of massABC deletion does not play a role in dauer suppression.

Because ΔmassA and ΔmassBC also lost twitching motility, we cannot formally exclude a contribution of pili-mediated motility without testing a pilus-deficient mutant in the dauer assay.

To test if massetolide plays a role in clearance of *P. lurida* cells in the *C. elegans* intestine, we visualized and quantified WT, *ΔgacA*, and *ΔmassBC P. lurida* cells in the intestine using FISH probes. FISH probe visualization revealed a higher intestinal bacterial load in animals fed *P. lurida ΔgacA* or *ΔmassBC* than in those fed the wild-type strain (**Supplemental Fig. 4**). These results suggest that GacS/GacA-regulated massetolide production is required for efficient clearance of *P. lurida* from the host intestine, as loss of either massetolide production or its upstream regulator resulted in increased accumulation of bacteria within the intestine. A similar phenotype was also observed in our collaborators’ co-submitted study(cit) in which *P. lurida*-derived massetolide promotes host-mediated clearance through TGF-β and serotonergic signaling and simultaneously renders bacteria susceptible to that clearance by modulating bacterial surface properties^28^.

### MassABC produces Massetolide F and Massetolide F restores dauer suppression in *massBC* mutants

Massetolides are cyclic lipopeptides produced by *Pseudomonas* species that contribute to bacterial surface behaviors and interspecies interactions, including swarming motility, biofilm formation, plant pathogen control, and protection of *C. elegans* against bacterial infection^11,18,22,35,37^. However, whether massetolides directly impact animal physiology in the context of reduced insulin or TGF-β signaling is unknown. Because closely related massetolide isoforms (A-L) can be produced by the same bacterial species^38^, we aimed to determine which massetolide isoform is predominantly produced by *P. lurida* MYb11 and whether this molecule is sufficient to restore dauer suppression in massetolide deficient mutants. To address this, we identified and purified massetolide isoform(s) produced by *P. lurida* MYb11 under the growth conditions we use in our dauer development assays. A comparative mass spectrometry analyses of the extract of the *Pseudomonas lurida* wild type versus the two mutant strains *ΔmassA* and *ΔmassBC* with a protonated molecular ion at m/z 1126.6889 (calcd for C54H96N9O16+, m/z 1126.6970) present only in the wild type suggesting the probable target compound (**Fig. 3A and 3B**). This compound was isolated from the extract of the *P. lurida* wild type by preparative HPLC (**Fig. 3C**) and was characterized by NMR spectroscopy (**Supplemental Fig. 5**). The NMR data matched that of the massetolide F reported in the literature^38^ and the mass spectrometry data also in agreement with it. We conclude that the *P. lurida massABC* biosynthetic cluster produces Massetolide F under the growth conditions we used. We note that the same cluster can produce Massetolide E which differs from Massetolide F by a single methylene in the peptide core, owing to the relaxed substrate specificity of the adenylation domains and that Massetolide E has also been found to be produced by *P. lurida* but we did not detect significant amounts of Massetolide E using our growth conditions^11,39^.

**Figure 3:**
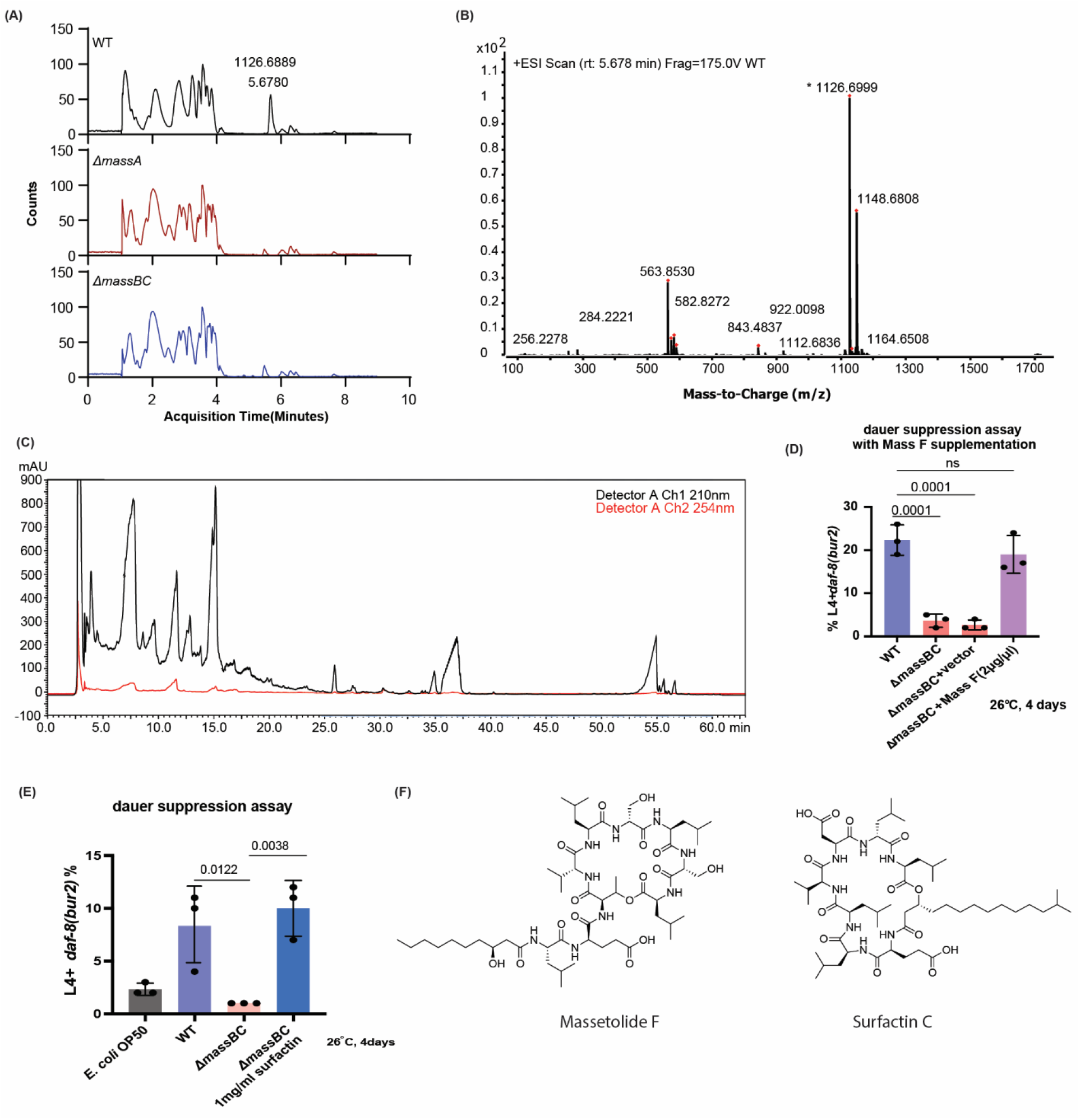
*P. lurida* MYb11 *massA* and *massBC* are required for the synthesis of Massetolide F. **(A)** LC-MS TICs of *P. lurida* WT, *ΔmassA* (NOBb1018), and *ΔmassBC* (NOBb1023) extract. The WT extract has a peak at 5.6780 minutes with an m/z of 1126.6889 that does not appear in either mutant strain. **(B)** Mass spectra of the feature at retention time 5.6780 minutes. Daughter ion fragmentation is constituent with Massetolide. **(C)** HPLC of WT MYb11. The peak at approximately 35.0 to 36 minutes was collected for NMR and supplemention studies. **(D)** Exogenous Massetolide F supplementation is sufficient to promote *daf-8*(*bur2*) animals to develop. *daf-8*(*bur2*) animals were incubated on WT MYb11 and *ΔmassBC*(NOBb1023) with either vector (10% DMSO) or Massetolide F dissolved in 10% DMSO (2µg/µL) for 4 days at 26° C. The number of L4+ animals was counted, and percentage of the total population is reported. Statistical significance was determined by ordinary one-way ANOVA followed by Sidak’s multiple comparisons testing. Error bars indicate s.d. from the mean. Error bars indicate s.d. from the mean. Exact p values are shown on the graph. **(E)** Exogenous surfactin supplementation is sufficient to promote *daf-8*(*bur2*) animals to develop. *daf-8*(*bur2*) animals were incubated on WT MYb11 and *ΔmassBC*(NOBb1018) with either vector (10% ethanol) or surfactin dissolved in 10% ethanol (1µg/µL) for 4 days at 26° C. The number of L4+ animals was counted and percentage of the total population is reported. Statistical significance was determined by ordinary one-way ANOVA followed by Sidak’s multiple comparisons testing. Error bars indicate s.d. from the mean. Error bars indicate s.d. from the mean. Exact p values are shown on the graph. **(F)** the chemical structures of Massetolide F compared to Surfactin C.

We next tested whether HPLC purified massetolide F could restore dauer suppressive activity to *ΔmassBC*. To test this, we supplemented *P. lurida* MYb11 Δ*massBC* bacterial cultures with purified massetolide F suspended in 100% DMSO at a final concentration of 2 µg/µL and quantified dauer arrest in *daf-8(bur2)* animals at 26 °C. We found that supplementation with massetolide F restored the ability of *P. lurida* MYb11 Δ*massBC* to suppress dauer arrest and promote development in *daf-8(bur2)* animals (**Fig. 3D**). These results demonstrate that massetolide F is sufficient to restore the dauer suppressive activity of *P. lurida* MYB11 Δ*massBC*. To determine whether dauer suppression is a general property of biosurfactants or cyclic lipopeptides, we also tested surfactin, a biosurfactant and cyclic lipopeptide produced by some strains of *Bacillus subtilis*^40^, and daptomycin, a cyclic lipopeptide antibiotic produced by *Streptomyces roseosporus*^41^. Similar to massetolide F, surfactin suppressed dauer formation and restored developmental progression in *daf-8(bur2)* animals fed the Δ*massBC* mutant (**Fig. 3E**). In contrast, daptomycin did not alter dauer arrest in *daf-8(bur2)* animals (**Supplemental Fig. 6**). These findings suggest that biosurfactants such as massetolide F and surfactin can promote development in animals with impaired TGF-β signaling, but that this activity is not a general property of all bacterially produced cyclic lipopeptides (**Fig. 3F**). These results reveal a potentially broader role for bacterial cyclic lipopeptides in regulating host development.

### Massetolide F alters dauer associated transcriptional programs in host animals

To determine how massetolide F affects host gene expression in animals with impaired TGF-β signaling, we sequenced the transcriptomes of *daf-8(bur2)* animals fed either wild type *P. lurida* MYb11 or *P. lurida* MYb11 Δ*massBC*. We first ranked genes by differential expression and focused on the most upregulated and downregulated genes (top 30) by massetolide F (**Fig. 4A; Supplementary Table 1**). GO enrichment analysis identified biological changes associated with the differentially expressed genes (**Fig. 4B**), while functional classification grouped these genes into broad biological categories (**Fig. 4C**). From these genes, we identified candidates with reported or potential connections to dauer biology (**Fig. 4D, Supplementary Table 3**). Among genes most upregulated by massetolide F, *asm-3* and *dod-21* were associated with dauer relevant signaling outputs. *asm-3* has been linked to the insulin signaling pathway, and higher *asm-3* expression is reported consistent with reduced dauer arrest^42^. *dod-21* is a DAF-16 associated output gene and may reflect altered insulin signaling or DAF-16 dependent transcriptional output^43,44^. Unlike *asm-3*, loss of *dod-21* is not known to affect dauer formation or exit. In contrast, several genes associated with dauer formation physiology were downregulated by massetolide F. These included *hsp-12.6* (encodes a small heat shock protein associated with dauer and stress physiology), *oac-39* (encodes an O acyltransferase linked to dauer associated maradolipid synthesis), and *scl-12* and *scl-13* (encode SCP like lipid binding proteins implicated in cholesterol handling during dauer formation and recovery) ^45–47^. Thus, bacterial massetolide F reduced the expression of several dauer formation related genes that are known to be downstream of the insulin and TGF-β signaling pathways. These findings are consistent with Massetolide F antagonizing the pathways that regulate dauer formation in *C. elegans*.

**Figure 4.**
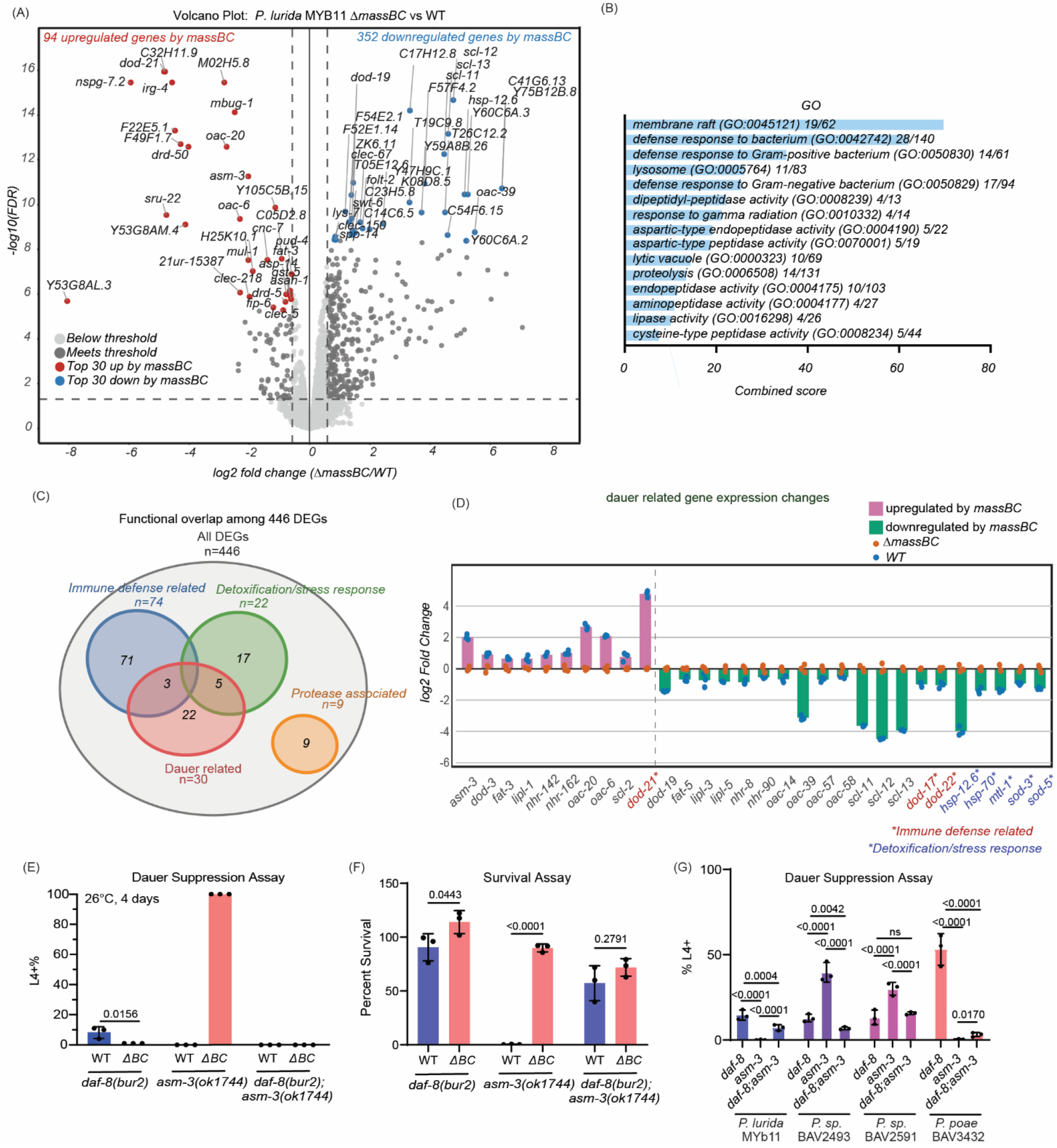
The *P. lurida massBC* biosynthetic cluster regulates host gene expression and promotes development through *asm-3*. **(A)** Volcano plot showing differential transcript abundance in *C. elegans* fed *P. lurida* MYB11 Δ*massBC* versus WT. Red dots indicate transcripts upregulated by *massBC*, blue dots indicate transcripts downregulated by *massBC*, dark gray dots indicate significant transcripts outside the top 30 in each category, and light gray dots indicate nonsignificant transcripts. Dashed lines indicate the significance thresholds of |Fold Change| >=1.5 and Adjusted P-Value <=0.05. **(B)** Gene Ontology (GO) enrichment analysis of the 446 DEGs. Enriched terms are ranked by combined score. Fractions following each term indicate the number of DEGs assigned to that term relative to the total number of genes associated with the corresponding GO category. **(C)** Functional overlap among the 446 differentially expressed genes, including genes associated with immune defense, detoxification or stress responses, dauer development, and protease activity. **(D)** Expression changes in reported dauer-related genes in animals fed *P. lurida* WT or Δ*massBC*. Magenta bars indicate genes upregulated by *massBC*, and green bars indicate genes downregulated by *massBC*. Orange and blue dots represent individual biological replicates from animals fed Δ*massBC* and WT bacteria, respectively. Red and blue asterisks indicate genes also associated with immune defense and detoxification or stress responses, respectively. Detailed information of these gene listed here are in Supplementary Table 3. **(E)** Dauer suppression assays showing the percentage of *daf-8(bur2)*, *asm-3(ok1744)*, and *daf-8(bur2); asm-3(ok1744) double mutant* animals that developed to the L4 stage or later after 4 days at 26°C on *E. coli* OP50, *P. lurida* WT, or *P. lurida* Δ*massBC*. Dots represent three independent biological replicates. Exact *P* values are shown for the indicated comparisons. **(F)** *massBC* are required for *asm-3*(*ok1744*) survival on *P. lurida*. Animals were incubated at 26 °C for 4 days. The percentage of animals that survived were calculated by normalizing to the total number of animals that survived on *E. coli* OP50 control plates (see SFigure 7). Experiments were performed in triplicate with approximately 500 animals per replicate. Statistical significance was determined by 2way ANOVA followed by Sidak’s multiple comparisons testing. Error bars indicate s.d. from the mean. Exact p values are shown on the graph; ns, not significant. **(G)** Dauer suppression assays showing the percentage of *daf-8(bur2)*, *asm-3(ok1744)*, and *daf-8(bur2); asm-3(ok1744)* double mutant animals that developed to the L4 stage or later after 5 days at 26°C on *E. coli* OP50, *P. lurida, Pseudomonas sp.* BAV2493*, Pseudomonas sp. BAV2591,* and *a* BAV3432. Animals were incubated at 26 °C for 5 days. The percentage of animals that survived were calculated by normalizing to the total number of animals that survived on *E. coli* OP50 control plates. Experiments were performed in triplicate with approximately 500 animals per replicate. To account for unequal variance, statistical significance was determined by transforming the percentages^87^ and modeling the data using beta regression^88^ followed by assessing the tests with emmeans and correcting for multiple testing using a Tukey’s HSD^89^. Error bars indicate s.d. from the mean. Exact p values are shown on the graph.

WormEnrichr analysis of the complete 446 differentially expressed genes (absolute fold change > 1.5 and adjusted P value <= 0.05) in *daf-8(bur2)* animals identified enrichment for defense response, proteolysis, peptidase activity, membrane raft, and lysosome associated terms (**Fig. 4B**). These results suggest that in addition to genes involved in dauer development and formation, massetolide F may also affect the expression of host immune and proteolytic programs. We therefore asked which massetolide F regulated genes were associated with host immune defense or stress response, and whether these genes overlapped with dauer related genes. Based on the RNA-seq data, we observed overlap between dauer related genes and immune defense or stress response gene clusters (**Fig. 4C and D, Supplementary Table 3**). These findings are consistent with the known changes in immune and antimicrobioal peptide expression that occur during dauer development and are regulated by TGF-β and insulin-like signaling. Lastly, we also performed the same RNA seq comparison in wild type N2 animals to test if the effects of Massetolide F on gene expression were specific to our insulin signaling or TGF-β mutant models or if Massetolide F also altered the physiology of wild-type animals. N2 animals also exhibited differential gene expression when fed *P. lurida* WT compared with the *ΔmassBC* mutant, indicating that the host transcriptional response associated with massetolide F is not restricted to animals with impaired dauer signaling (**Supplementary Table 2, Supplementary Figure 7**). Together, these results indicate that exposure to massetolide F is associated with broad changes in host gene expression in both *daf-8(bur2)* and WT animals.

### The sphingomyelinase, *asm-3*, is required for Massetolide F to promote development of TGF-β mutants

*asm-3* was previously reported to alter dauer formation and is differentially activated by the microbiota^42,48–50^. In addition, *asm-3* was the most upregulated gene that we identified to be regulated by massetolide F that is also known to be involved in regulating dauer formation. Thus, we further investigated whether host *asm-3* is required for Massetolide F to suppress dauer formation. *asm-3* encodes an acid sphingomyelinase that hydrolyzes sphingomyelin to generate ceramide, thereby contributing to sphingolipid turnover and the regulation of cellular membrane composition and signaling^51,52^. When fed *P. lurida* WT, *daf-8(bur2); asm-3(ok1744)* double mutants remained developmentally arrested (**Fig. 4E**), indicating that *asm-3* is required for *P. lurida* to suppress dauer formation in our mutant models. To test if *asm-3* single mutants are themselves *daf-c* under the same conditions we grew *asm-3(ok1744)* single mutants on *E. coli* OP50 under identical conditions and found that 100% of *asm-3(ok1744)* single mutants grew to adulthood, suggesting that loss of *asm-3* does not cause dauer development on its own (**Fig 4E**). Surprisingly, however, we found that when we placed *asm-3(ok1744)* mutant embryos on wild-type *P. lurida* nearly 100% of embryos died embryonically and that this embryonic lethality was dependent on the production of Massetolide F (**Fig. 4F, SFig. 8**). This impact of Massetolide F on *asm-3(ok1744)* single mutants occurs before animals feed on *P. lurida*, suggesting that Massetolide F can impact *asm-3* mutant *C. elegans* independently of any change in nutrition or intestinal colonization (embryos have not yet fed on *P. lurida*). In addition to these unexpected findings, we found that *P. lurida* was no longer toxic to *daf-8(bur2); asm-3(ok1744)* double mutant animals (**Fig. 4F**). This finding indicates that the loss of *asm-3* is not absolutely required for survival in the presence of Massetolide F, but rather that *asm-3* and *daf-8* likely antagonistically regulate animal development. Collectively, these results indicate that Massetolide F induces the expression of *asm-3* and that Massetolide F induced dauer suppression requires *asm-3*. To determine if *asm-3* is broadly required for *Pseudomonas*-mediated suppression of *daf-8*, we placed *daf-8*(*bur2*), *asm-3*(*ok1744*), and *daf-8*(*bur2*);*asm-3*(*ok1744*) embryos on the panel of *Pseudomonas* species that suppressed dauer (see figure 1E). We found fewer *daf-8*(*bur2*);*asm-3*(*ok1744*) animals developed beyond dauer arrest than *daf-8*(*bur2*) animals. These results indicate that *asm-3* is partially required for suppression of *daf-8(bur2)* by different *Pseudomonas* species, suggesting either 1) Masetolide F is conserved in these species or 2) *asm-3* plays a broader role in dauer signaling.

## Discussion

Microbiome-derived metabolites can regulate host physiology, but linking defined bacterial genes and metabolites to specific host phenotypes remains a major challenge. Here, we performed a large-scale screen using *C. elegans* dauer formation as a genetically sensitized readout to identify bacterial mechanisms that promote development when TGF-β or insulin signaling is impaired. This approach led to the discovery that Massetolide F, a cyclic lipopeptide synthesized in part by *P. lurida* MassABC, induces host *asm-3* expression and thereby promotes development of TGF-β mutant animals (*daf-8(bur2)*) (**Fig. 5**). In conjugation with our collaborators, we found Massetolide F is required for *P. lurida* motility and bacterial clearance from the *C. elegans* intestine. We also found that the structurally distinct bacterial cyclic lipopeptide, surfactin, is sufficient to suppress *daf-8(bur2)* dauer arrest, similarly to Massetolide F. Thus, the suppression of dauer is not specific to Massetolide F and might be mediated by a subset of cyclic lipopeptides which are produced by diverse species. Overall, these findings demonstrate that a subset of bacterial cyclic lipopeptides either directly or indirectly leads to the suppression of TGF-β and insulin mediated developmental arrest.

**Figure 5:**
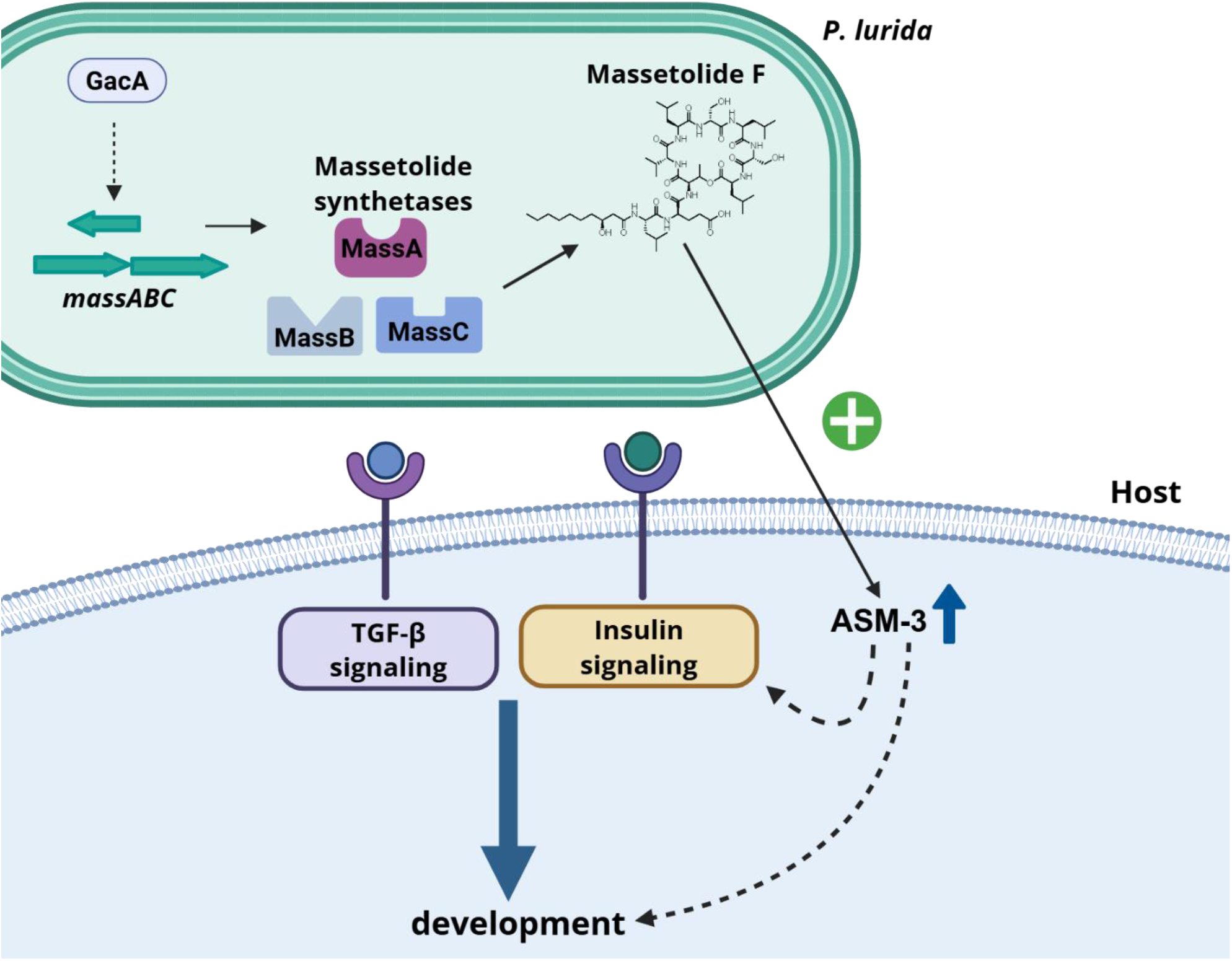
Model for how Massetolide F that is synthesized by *P. lurida* and regulated by the *gacA* regulatory pathway suppresses TGF-β and insulin signaling mutants by activating *asm-3* in *C. elegans*.

In the last 10-20 years, *C. elegans* has been adapted to screen for novel host-microbe interactions and dietary molecules that regulate animal physiology ^8,13,24,27,53,54^. Here, we further build on the benefits of this model organism (well established genetic models, easily collectable germ-free animals) to identify small bacterial molecules that promote *C. elegans* development using dauer arrest as a gross morphological readout. To the best of our knowledge, this work is the first to report the successful identification of bacterial strains and genes that lead to suppression of TGF-β and insulin mutant dauer arrest. While this work focuses on a single possible mechanistic example of suppression of *daf-c* mutant animals, we propose that our *C. elegans*-based screening platform could be used to identify additional, unknown mechanisms by which bacterial secondary metabolites alter the regulation of animal development across a nearly any molecular pathway that is important for animal development. Similarly, the same general approach used here should be adaptable to identify bacterial species and molecules that regulate any easily assayed change in *C. elegans* biology including GFP expression, survival, or fertility. Thus, we expect that similar approaches using *C. elegans*-based high-throughput screens might serve as a critical future tool for linking bacterial strains, genes, and metabolites to specific changes in animal physiology.

Previous work has established that Massetolides and related cyclic lipopeptides influence bacterial surface motility, biofilm formation, antimicrobial activity, plant-microbe interactions, and microbial competition^18,22,23^. Specifically, previous work suggested that massetolide in *P. lurida* has antimicrobial activity^11,12^. Our work suggests new mechanisms for how cyclic lipopeptides benefit host biology. First, we found that loss of Massetolide F resulted in higher intestinal bacterial burden in the host, suggesting that Massetolide F may protect the host from bacterial accumulation in the intestine. Bacterial burden in the *C. elegans* intestine is well-established to play a role in lifespan and susceptibility to pathogens^55–57^. Thus, cyclic lipopeptide-mediated control of the intestinal bacterial load may help maintain intestinal homeostasis and promote host health. Second, we found *P. lurida massABC* increased *asm-3* expression **(Fig. 4D)** and that massetolide F-mediated suppression of *daf-8(bur2)* dauer requires *asm-3* **(Fig. 4E)**. ASM-3 is an acid sphingomyelinase that converts sphingomyelin to ceramide^51^ and has been reported to positively regulate animal development via DAF-2 insulin signaling^42^. However, how Massetolide F drives *asm-3* expression and specifically how *asm-3* drives *daf-2* signaling remains unknown. Several studies have previously established that diverse bacterial species increase host *asm-3* expression and that vitamin B12 may be required in one of these models ^50,58^. Previous studies have also established that bacterial sphingolipids can enter host sphingolipid pathways and alter ceramide pools, suggesting a model by which bacterial sphingolipids activate host *asm-3*^48,59–61^. We found that Massetolide F, rather than B12 or sphingolipids, leads to an increase in *asm-3* expression; however, Massetolide F may drive *asm-3* activation via an indirect mechanism. Future studies will likely be critical in answering these outstanding mechanistic questions.

We found the cyclic lipopeptide surfactin is also sufficient to suppress *daf-8(bur2)* dauer arrest, suggesting a broader subset of cyclic lipopeptides may lead to the activation of *asm-3* mediated animal development. To our knowledge, surfactin has not previously been reported to regulate dauer formation or TGF-β signaling in *C. elegans*. However, surfactin has been reported to improve insulin resistance through PI3K/Akt- and AMPK-associated mechanisms in mammalian cell and mouse models^62^ and to alter TGF-β expression during wound healing and scar formation^63^. These findings raise the exciting possibility that cyclic lipopeptides like Massetolide F and sufactin might generally promote increased TGF-β and insulin signaling across diverse animal species ranging from invertebrates to mammals. Whether surfactin induced insulin signaling and TGF-β expression relies on ASM (mammalian ortholog of *asm-3*) in mammalian models remains unknown. The lack of activity observed with daptomycin further indicates that dauer suppression is not a general property of bacterial cyclic lipopeptides. We hypothesize that specific structural or biophysical properties shared by biosurfactants such as Massetolide F and surfactin may contribute to their ability to promote development under conditions of impaired TGF-β signaling. Further studies will be required to determine whether surfactin and Massetolide F act through shared host targets or through distinct mechanisms. Also, future studies comparing our findings in *C. elegans* with previous findings in mouse models will be critical in determining how cyclic lipopeptides might mechanistically alter animal physiology across species.

In summary, this study identifies Massetolide F and surfactin as bacterial cyclic lipopeptides that promotes development in animals with impaired dauer regulatory signaling. By linking GacS/GacA regulation, the *massABC* biosynthetic cluster, purified Massetolide F, host ceramide metabolism, and development outcomes, this work provides a mechanistic example of how a bacterial biosurfactant can regulate animal physiology. More broadly, these findings suggest that microbial molecules traditionally studied for surface activity or antimicrobial function may also include host-active metabolites with potential relevance to conserved disease-associated pathways. Lastly, these findings further support the usefulness of the *Caenorhabditis elegans* model system to study host-microbe interactions and particularly to identify previously unknown bioactive gene clusters and metabolites and influence animal physiology.

## Methods

Research with *C. elegans* does not require IACUC approval.

### *C. elegans* strains, growth media, and conditions

*C. elegans* strains were maintained at 15 °C or 20 °C on Nematode Growth Medium (NGM) agar plates seeded with *E. coli* OP50 (CGC). NGM agar contained 3 g/L NaCl (VWR, BDH9286), 17 g/L agar (Sigma-Aldrich, A1296), 2.5 g/L peptone (Bacto, 211677), 25 mM KPO₄, pH 6.0 (Sigma-Aldrich, P0662 and P3786), 5 mg/L cholesterol (Sigma-Aldrich, C8503), 1 mM MgSO₄ (VWR, 0662), and 1 mM CaCl₂ (VWR, 0556). All *C. elegans* strains used in this study are listed in **Supplementary Table 4**. To generate homozygous double mutant strains, genetic crosses were performed between animals carrying the two mutations of interest. Young L4 hermaphrodites carrying one mutation were mated with males homozygous for the second mutation on NGM agar plates for 24–48 hours. Heterozygous F1 hermaphrodites carrying both mutations were transferred to separate NGM plates and allowed to self-fertilize. Individual F2 progeny were then isolated and genotyped by Sanger sequencing or PCR-based genotyping to identify animals homozygous for both mutations.

### Egg preparation protocol

Adult *C. elegans* were washed from NGM agar plates using M9 buffer containing 3 g/L KH₂PO₄ (Sigma-Aldrich, P0662), 6 g/L Na₂HPO₄ (Sigma-Aldrich, P0662, P3786), 5 g/L NaCl (VWR, BDH9286), and 1 mM MgSO₄ (VWR, 0662). Animals were lysed in egg preparation solution containing 30% sodium hypochlorite and 1 M NaOH. Lysis was monitored using a dissecting microscope for approximately 5 to 10 min, until most adult carcasses were degraded and embryos remained intact. Embryos were then washed two to three times with M9 buffer to remove residual egg preparation solution.

### Bacterial strains, growth media, and conditions

All bacterial strains were grown on Lennox LB agar or in Lennox LB broth (Invitrogen 12780-029, Sigma Aldrich A1296) at 30 °C or 37 °C unless specified otherwise. All bacterial strains used in this study are listed in **Supplementary Table 4**. Liquid cultures were grown in ≤5 mL Lennox LB in 18 mm glass test tubes at 30 °C with shaking at 220 rpm. Plasmid selection in *E. coli* was performed on Lennox LB containing kanamycin (50 µg/mL; Sigma-Aldrich, K1377) or tetracycline (10 µg/mL; Sigma-Aldrich, A0166), depending on the plasmid antibiotic resistance marker. *P. lurida* MYB11 strains carrying pBBR1 derivatives for complementation were grown in Lennox LB containing kanamycin (25 µg/mL; Sigma-Aldrich, K1377). Vogel-Bonner minimal medium (VBMM) plates containing tetracycline (10 µg/mL) were used to select *P. lurida* transconjugants and counter select against the *E. coli* donor strain ^64,65^. VBMM plates were prepared from 10× VBMM salts containing 2 g MgSO₄·7H₂O, 20 g citric acid, 100 g K₂HPO₄, and 35 g NaNH₄HPO₄·4H₂O per liter, adjusted to pH 7.0 and filter sterilized ^65^. To prepare VBMM agar plates, 100 mL of 10× VBMM salts was added to 900 mL of sterile molten agar containing 15 g agar per final liter, cooled, and supplemented with tetracycline to desired concentration.

### Bacterial gene deletion and complementation

Bacterial gene deletion mutants were generated using an allelic exchange-based strategy modified from previously described methods^64,66^. Briefly, upstream and downstream homology arms for each target gene or operon were amplified from *Pseudomonas lurida* MYB11 genomic DNA by touchdown PCR using Q5 DNA polymerase (New England Biolabs, M0492) and oligonucleotides from Integrated DNA Technologies (Coralville, IA) listed in **Supplementary Table 5**. PCR fragments were assembled into linearized pEX18Tc (digested with EcoRI and HindIII) using NEBuilder HiFi DNA Assembly Master Mix (New England Biolabs, E2621).

Assembled plasmids were transformed into *E. coli* DH5α competent cells (New England Biolabs, C2988J) according to the manufacturer’s instructions for propagation and sequence verification. All PCR amplified regions were confirmed by DNA sequencing at Genewiz from Azenta Life Sciences (South Plainfield, NJ). Verified deletion plasmids were then transformed into *E. coli* S17-1 λpir (ATCC, BAA-2428) and transferred into *P. lurida* MYB11 by conjugation. Transconjugants were selected on VBMM plates containing tetracycline (10 µg/mL), and deletion mutants were isolated by sucrose counterselection. Candidate deletion mutants were screened by PCR using primers flanking the target region and confirmed by Sanger sequencing. All bacterial mutants generated in this study are listed in **Supplementary Table 4**, and plasmids are listed in **Supplementary Table 6**.

For complementation, each target gene was amplified together with its native promoter region from *P. lurida* MYB11 genomic DNA. The resulting fragment was assembled into the broad host range plasmid pBBR1^67,68^ linearized with EcoRI and HindIII and using NEBuilder HiFi DNA Assembly Master Mix (New England Biolabs, E2621). Complementation plasmids were verified by Sanger sequencing in *E. coli* DH5α and introduced into the corresponding *P. lurida* deletion mutant strains by conjugation as described above. Complemented strains were maintained on LB or NGM containing kanamycin (25 µg/mL).

The P*_massA_*’-‘*lacZ* reporter was cloned by amplifying truncated *lacZ* using primers KN668 and KN669 (listed in Supplemental Table 5) from pUC18-mini-Tn7T-GM-lacZ^69^. pUC18-mini-Tn7T-Gm-lacZ was a gift from Herbert Schweizer (Addgene plasmid # 63120 ; http://n2t.net/addgene:63120 ; RRID:Addgene_63120). The promoter region and first 180 bps of MYb11 *massA* was amplified using the primers KN666 and KN667 (listed in Supplemental Table 5). The amplicons were assembled into BBR1-MCS5^67^ digested with AgeI and Nsil (NEB#R3552S and #R3127S respectively) by isothermal assembly (New England Biolabs, E2621) generating pNOB05. The promoter-*lacZ* fusion was then amplified from pNOB05 using primers KN711 and KN712 and assembled in pUC18t-mini-TN7t-Gm digested with BamHI and HindIII generating pNOB12. The reporter was electroporated in *P. lurida* MYb11 strains as previously described^70^. Reporter construct integration was validated with whole genome sequencing at SeqCenter (Pittsburgh, PA).

### Dauer suppression assay

Dauer suppression assays were performed using synchronized *C. elegans* embryos generated by the egg preparation protocol described above. Bacterial strains of interest were grown overnight in Lennox LB at 30 °C with shaking, and cultures were seeded onto 60 mm NGM plates with 500 µL culture, 90 mm NGM plates with 1 mL culture, or individual wells of 12 well NGM plates with 200 µL culture. Bacterial strains carrying plasmids that required antibiotic selection were seeded on NGM supplemented with the appropriate antibiotics at the indicated concentrations. Seeded plates were dried in a laminar flow hood before embryos were added. Approximately 100 to 200 embryos were added to each well, or approximately 500 embryos were added to each plate. Plates were incubated at the dauer forming temperature reported for each *C. elegans* strain from Caenorhabditis Genetics Center (CGC), for example 26 °C for 4 days for *daf-8(bur2)* and *daf-1(m40)*. Animal developmental stage was then scored using a dissecting microscope. Animals that remained as dauers were scored based on dauer morphology, including developmental arrest and a thin, radially constricted body shape, whereas animals that developed to the L4 stage or adulthood were scored as developmentally rescued. Dauer suppression was calculated as the percentage of animals that developed to the L4 stage or later. Each condition was tested with three biological replicate plates, and bacterial strains or treatments that altered dauer arrest were retested in independent assays. For Massetolide F specific dauer suppression assays, 35 mm × 10 mm NGM plates (Millipore Sigma, CLS430165) were used because of the limited amount of purified Massetolide F available. For these assays, 50 to 60 embryos were added to each plate, with three biological replicate plates per condition. Other cyclic lipopeptides used in dauer suppression assay are surfactin (Sigma-Aldrich, S3523) and daptomycin (Thermo Scientific Chemicals, AC461375000).

### Screen of wild bacterial isolates for *C. elegans* development

Compost samples were collected from the Grand Rapids, MI area and resuspended in M9 buffer. The suspensions were diluted, filtered, and plated on LB agar to isolate individual bacterial colonies. 4,000 single colonies were picked and grown overnight at 30 °C, and 200 µL of each overnight culture was seeded into individual wells of NGM 12 well plates. In parallel, bacterial cultures were saved in individual wells of 96 well plates to generate a wild bacterial isolate library. Seeded NGM plates were dried in a laminar flow hood before approximately 100 to 200 germ free *C. elegans daf-1(m40)* embryos were added to each well. Plates were incubated at 26 °C for 4 days, and animal development was scored by eye using a dissecting microscope. The percentage of animals that developed to the L4 stage or later was used as the primary readout. Bacterial isolates that suppressed dauer formation and promoted *C. elegans* development in the primary screen were retested on 90 mm NGM plates for confirmation.

### Microscopy

*C. elegans* were washed three times with M9 buffer before imaging to remove excess bacteria.Animals were paralyzed in tetramisole at 10 µg/mL and incubated for approximately 5 min before imaging. Images were taken using a Leica DM6 B upright microscope equipped with a Leica DFC9000 GT camera with 4.2-megapixel resolution (2048 × 2048 pixels), using 4x or 10x objective lenses and a 6.5 µm × 6.5 µm pixel size. Leica Application Suite X version 3.7.5.24914 was used for image acquisition. Images were assembled and analyzed using Adobe Illustrator 2024 version 28.4.1, 64 bit.

### Transposon mutagenesis and screening

A *P. lurida* MYB11 transposon mutant library was generated using the EZ-Tn5 Tnp Transposome kit (Biosearch Technologies, TSM99K2) according to the manufacturer’s protocol. Electrocompetent *P. lurida* MYB11 cells were prepared from concentrated cultures by washing four times with ice cold 10% glycerol. For each electroporation, 50 µL of electrocompetent cells was mixed with 1 µL of transposome and pulsed in a 0.2 cm gap cuvette using a Gene Pulser Xcell electroporation system (Bio-Rad Laboratories) with manual settings used for *Pseudomonas aeruginosa*. Cells were recovered in 1 mL LB at 30°C for 1 h with shaking, pelleted, resuspended in 100 µL LB, and plated on LB agar containing kanamycin (25 µg/mL). After overnight growth at 30°C, individual kanamycin resistant colonies were picked into 96 well plates containing LB with kanamycin (25 µg/mL) and stored at -80 °C. For screening, individual transposon mutants were grown overnight at 30 °C in 1.5 mL LB with kanamycin (25 µg/mL) in 18 mm glass test tubes. Cultures were seeded into wells of 12 well NGM plates containing kanamycin (25 µg/mL) and dried in a laminar flow hood. Approximately 100 to 200 germ free *C. elegans daf-8(bur2)* embryos were added to each well, and plates were incubated at 26 °C for 4 days. Mutants that failed to suppress dauer arrest or promote animal development were selected as candidate hits and retested on 90 mm NGM plates for confirmation.

### Bacterial genomic DNA extraction and whole genome sequencing

Single bacterial colonies were grown overnight in 15 mL liquid culture at 30 °C. Cultures were centrifuged at 17,000 × g for 5 min, and genomic DNA was isolated using the Quick-DNA Fungal/Bacterial Miniprep Kit (Zymo Research, D6005) according to the manufacturer’s instructions. Briefly, bacterial pellets were resuspended and transferred to ZR BashingBead lysis tubes containing BashingBead buffer. Cells were lysed by bead beating, and lysates were centrifuged to remove debris. The cleared supernatant was passed through a Zymo-Spin III-F filter, mixed with genomic lysis buffer, and loaded onto a Zymo-Spin IICR column. Bound DNA was washed with DNA pre-wash buffer and gDNA wash buffer, then eluted in deionized water. Genomic DNA concentration and purity were measured using a NanoDrop spectrophotometer, and DNA integrity was assessed by visualization on a 1% agarose gel.

Genomic DNA from bacterial isolates of interest was submitted for whole genome sequencing on an Illumina PE150 platform at SeqCenter (Pittsburgh, PA). FastQC version 0.12.1 was used to assess raw sequencing read quality^71^. Reads were trimmed using Trimmomatic version 0.39 to remove adapter sequences and low-quality reads with Q < 30^72^. Genome assembly, annotation, and variant calling were performed using the Bacterial and Viral Bioinformatics Resource Center (BV-BRC). Within BV-BRC, bacterial genomes were assembled using Unicycler with default parameters^73^. Genome annotation was performed using RASTtk^74^. For variant calling, reads were aligned to the reference genome using BWA-MEM^75^ and variants were identified using FreeBayes^76^ through the SNP Caller workflow.

### Sample Preparation for Proteomics

The cell pellets were homogenized on the Bead Ruptor Elite (Cat# 19-042E, Omni International) for 30 s in 4% SDS solution containing 1x HALT Protease (Cat# 78442, Thermo Fisher Scientific). Samples were sonicated and clarified via centrifugation and transferred to a Protein LoBind Eppendorf tube. Proteins were quantified using the Pierce BCA Protein Assay Kit (Cat# 23227, Thermo Fisher Scientific) and 100 µg of protein was aliquoted for digestion. Protein digestion utilized the S-Trap (Cat# CO2-Mini, Protifi) platform to remove any SDS prior to LCMS/MS analysis. Briefly proteins were reduced with Dithiothreitol (DTT) for 20 minutes, alkylated with Iodoacetamide (IAA) for 20 minutes and digested overnight with Trypsin/Lys-C (Cat# V5072, Promega) at a ratio of 50:1 (protein:enzyme (w/w)). Peptides were eluted off the S-Trap Column and dried down in a Genevac SpeedVac. Dried samples were then cleaned up with Affinisep PurePep-SPE mini spin columns (Cat# Spin-PurePep.S.50, Affinisep) and dried down before resuspension for LC-MS/MS analysis. Dried samples were resuspended in 50 µL 0.1% FA (LS118-1, Fisher Scientific) and diluted with 50 µL of 0.1% TFA (LS119-500, Fisher Scientific).

### Data-independent Acquisition (DIA) LC-MS/MS Proteomics

DIA analyses were performed on Orbitrap Eclipse coupled to Vanquish Neo system (Thermo Fisher Scientific) with a FAIMS Pro source (Thermo Fisher Scientific) located between the nanoESI source and the mass spectrometer. 2 μg of digested peptides were separated on a nano capillary column (20 cm × 75 μm I.D., 365 μm O.D., 1.7 μm C18, CoAnn Technologies, Washington, # HEB07502001718IWF) at 300 nL/min. Mobile phase A consisted of LC/MS grade H_2_O (LS118-500, Fisher Scientific), mobile phase B consisted of 20% LC/MS grade and H_2_O and 80% LC/MS grade acetonitrile (LS122500, Fisher Scientific), both containing 0.1% FA. The LC gradient was: 1% B to 24% B in 110 min, 85% B in 5 min, and 98% B for 5 min, with a total gradient length of 120 min. The column temperature was kept constant at 50 °C using a customized column heater (Phoenix S&T, Chadds Ford, PA). For FAIMS, the selected compensation voltage (CV) was applied (−40V, -55V, and -70V) throughout the LC-MS/MS runs. Full MS spectra were collected at 120,000 resolution (full width half-maximum; FWHM), and MS2 spectra at 30,000 resolutions (FMWH). Both the standard automatic gain control (AGC) target and the automatic maximum injection time were selected. A precursor range of 380-985 m/z was set for MS2 scans, and an isolation window of 50 m/z was chosen with a 1 m/z overlap for each scan cycle. 32% HCD collision energy was used for MS2 fragmentation.

DIA data was processed in Spectronaut (version 19, Biognosys, Switzerland) using direct DIA. Data was searched against the *P. lurida and P. proteolytica* proteomes as appropriate, including expected mutations and isoforms. The manufacturer’s default parameters were used. Briefly, trypsin/P was set as digestion enzyme and two missed cleavages were allowed. Cysteine carbamidomethylation was set as fixed modification, and methionine oxidation and protein N-terminus acetylation as variable modifications. Identification was performed using a 1% q-value cutoff on precursor and protein levels. Both peptide precursors and protein false discovery rate (FDR) were controlled at 1%. Ion chromatograms of fragment ions were used for quantification. For each targeted ion, the area under the curve between the XIC peak boundaries was calculated.

### RNA sequencing and data analysis

Germ free *C. elegans* embryos were placed onto standard 90 mm NGM plates seeded with either *P. lurida* MYB11 WT or *P. lurida* MYB11 Δ*massBC*. Animals were maintained at 20 °C until adulthood, approximately 3 days. Each condition was performed in biological triplicate, with approximately 500 animals per replicate. Adult animals were collected and washed three times with M9 buffer to remove residual bacteria. Worm pellets were transferred to 750 µL ZymoBIOMICS DNA/RNA Shield, and total RNA was extracted using the ZymoBIOMICS RNA Miniprep Kit (Zymo Research Corporation) according to the manufacturer’s instructions. Briefly, samples in DNA/RNA Shield were homogenized in ZR BashingBead lysis tubes to disrupt worm tissue and associated bacterial material. Lysates were clarified by centrifugation, and the cleared supernatant was mixed with DNA/RNA lysis buffer before RNA purification. RNA was bound to a Zymo spin column, treated with DNase I on column to reduce genomic DNA contamination, washed with the provided buffers, and eluted in DNase/RNase free water. RNA concentration and purity were assessed using a NanoDrop spectrophotometer and Qubit fluorometer. Purified RNA samples were shipped on dry ice to Plasmidsaurus Inc. (South San Francisco, CA) for sequencing library preparation and RNA sequencing on an Illumina NovaSeq platform.

RNA sequencing data were processed using the Plasmidsaurus RNA seq analysis pipeline^77^. Briefly, FASTQ files were generated using BCL Convert v4.3.6 and fqtk v0.3.1, and reads were filtered with fastp v0.24.0 to remove poly-X tails and low-quality 3′ ends, using a minimum Phred quality score of 15 and minimum read length of 50 bp^78^. Filtered reads were aligned to the *C. elegans* reference genome *C. elegans*-WBcel235 2023-1-06_11.13.48_v14 using STAR v2.7^79^. BAM files were sorted with samtools v1.21, and PCR or optical duplicates were removed by UMI based deduplication using UMICollapse v1.1.0^80^. Gene expression was quantified using featureCounts from the Subread package v2.1.1 with strand specific counting and exon and 3′ UTR features grouped by gene_id^81^. Quality control, sample correlation, and principal component analysis were used to assess read quality, replicate consistency, and potential batch effects. Differential expression was analyzed using edgePython v0.2.5, and genes with an FDR corrected P value less than 0.05 were considered differentially expressed. For downstream analyses, differentially expressed genes were further filtered using absolute fold change greater than 1.5 and adjusted P value less than or equal to 0.05. Gene enrichment analysis was performed using WormEnrichr^82^.

### RNA FISH staining of intestinal bacteria

RNA fluorescence in situ hybridization (FISH) was performed to visualize alive bacteria in the *C. elegans* intestine using a protocol modified from previously described methods^83^. *C. elegans* animals were grown on the bacterial strains of interest at 20 °C until young L4. Young L4 animals were transferred to NGM plates supplemented with 50 µM 5-fluoro-2′-deoxyuridine (FUdR, Sigma-Aldrich, F0503) to prevent progeny production and seeded with the same bacterial diet. The animals were maintained with sufficient food for an additional 2 weeks after reaching the L4 stage. Worms were then washed from plates using M9 buffer containing 0.1% Tween 20 and transferred to 1.5 mL microcentrifuge tubes. Animals were pelleted by gentle centrifugation and washed three times with PBS containing 0.1% Tween 20 to remove external bacteria. Samples were fixed with acetone, protected from light, and stored at -20 °C until hybridization. Before probe hybridization, fixed animals were washed in hybridization buffer containing 900 mM NaCl, 20 mM Tris pH 7.5, and 0.01% SDS. General region of bacterial 16S rRNA was detected using a custom EUB338-CF610 probe ordered from LGC Biosearch Technologies, with the sequence 5′-GCTGCCTCCCGTAGGAGT-3′, a 5′ CAL Fluor Red 610 fluorophore, and a 3′ phosphate modification. The probe was added to hybridization buffer at a final concentration of 10 ng/µL, and samples were incubated overnight at 46 °C protected from light. After hybridization, animals were washed with wash buffer containing 900 mM NaCl, 20 mM Tris pH 7.5, 5 mM EDTA, and 0.01% SDS at 48 °C to remove unbound probe. Samples were then washed in PBS containing 0.1% Tween 20 and counterstained with DAPI to visualize host and bacterial DNA. After DAPI staining, animals were washed again in PBS containing 0.1% Tween 20, mounted on glass slides with antifade mounting medium, and imaged using fluorescence microscopy. Intestinal bacterial clearance was assessed based on CAL Fluor Red 610 signal localized within the intestinal lumen, with DAPI used as a nuclear and bacterial DNA counterstain.

### Massetolide Identification - Fermentation and extraction

Seed culture of *Pseudomonas lurida* was prepared in RPI Lennox Lysogeny Broth (LB) and incubated at 30°C for 24 h. This inoculum was used to seed six 2.8 L Fernbach flasks, each containing 1 L of LB medium (10 g/L casein digest peptone, 5 g/L yeast extract, and 5 g/L NaCl). The medium was adjusted to pH 7.0 prior to sterilization. Each flask was inoculated with 10 mL of the seed culture (1% v/v final concentration), resulting in a total working volume of 6 L. Fermentation was conducted at 30°C with orbital shaking at 150 rpm for 24 h. Following the growth phase, 40 g of pre-weighed Amberlite XAD-16 resin, contained in a mesh bag, was added to each flask. The flasks were agitated at room temperature for an additional 4 h to facilitate the adsorption of secondary metabolites. The resin bags were subsequently retrieved and rinsed with deionized water to remove residual fermentation broth. Bound compounds were eluted from the resin using methanol. The resulting methanolic extracts were pooled and concentrated to yield a crude extract for downstream chemical analysis.

The fermentation and extraction procedures described above were replicated for the *P. lurida* mutant strains, *ΔmassA* and *ΔmassBC*. For these experiments, the total culture volume was scaled to 2 L per strain (distributed across two 2.8 L Fernbach flasks). All parameters—including media composition, inoculation percentage (1% v/v), incubation conditions (30°C, 150 rpm), and the *in-situ* resin adsorption protocol—remained identical to the wild-type process to ensure experimental consistency for downstream comparative analysis.

### Massetolide Identification - Reagents and instruments

Chemicals were purchased from Sigma-Aldrich (St. Louis, MO, USA) unless otherwise specified. All the solvents used in extraction were of analytical-grade, while the solvents used for preparative HPLC and analytical UHPLC-MS were of HPLC-grade and Optima LC-MS grade, respectively, and supplied by Fisher Chemicals (Fair Lawn, NJ, USA). Prior to use, the solvents were filtered through a 0.45 μm polytetrafluoroethylene membrane. Deuterated solvent was purchased from Cambridge Isotopes Laboratory (Tewksbury, MA, USA). Preparative HPLC was performed on Shimadzu LC-20AP HPLC instruments with corresponding detectors, fraction collectors, and software. Ultra-high-performance liquid chromatograms (UHPLC) were obtained on an Agilent LC–MS system (Santa Clara, CA, USA) composed of an Agilent 1290 Infinity II UHPLC coupled to an Agilent 6545 ESI-qTOF-MS in positive mode using Kinetex C18 (2.6 µm, 3 × 75 mm) column eluted with 2 min isocratic elution of 90% A (A: 95 % H_2_O + 5 % ACN + 0.1% formic acid) followed by 5 min linear gradient elution to 100% B (95 % MeCN + 5 % H_2_O + 0.1% formic acid) with a flow rate of 0.4 mL/min. ESI conditions were set with the capillary temperature at 320 °C, source voltage at 3.5 kV and a sheath gas flow rate of 11 L/min. Ions detected in the full scan at an intensity above 1000 counts at 6 scans/s, with an isolation width of 1.3 ∼m/z, a maximum of 9 selected precursors per cycle and using ramped collision energy (5× m/z/100 + 10 eV). Purine C_5_H_4_N_4_ [M+H]^+^ ion (m/z 121.0508) and hexakis (1H,1H,3H-tetrafluoropropoxy)-phosphazene C_18_H_18_F_24_N_3_O_6_P_3_ [M+H]^+^ ion (m/z 922.009798) were used as internal lock masses. For high-resolution mass spectra (HR-ESIMS), ESI conditions were set with the capillary temperature at 320 °C, fragmentor voltage at 140 V, source voltage at 3.5 kV and a sheath gas flow rate of 11 L/min. Ions were detected in the full scan at an intensity above 1000 counts at 10 scans/s, with an isolation width of 1.3 ∼m/z. NMR data was recorded on a 600 MHz Varian/Agilent NMR Spectrometer ( 1 H: 600 MHz, 13C: 150 MHz) equipped with a 5 mm DB AUTOX PFG liquid N2 chilled Bruker Prodigy (^1^H/^19^F)-X broadband cryoprobe and a Bruker Avance NEO600 NMR system console, with automated tuning and matching (ATMA). MeOH-*d_3_* was used as the solvent for NMR. All NMR data analysis were manipulated using MestReNova NMR software, version 16.0.0 (Mestrelab Research, Santiego de Compostela, Spain).

### β-galactosidase Assay

WT MYb11 (NOBb263) and MYb11 *ΔgacA* (NOBb441) were grown overnight in LB at 30°C, 220 rpm. 1 mL of each overnight culture was pelleted and resuspended in Z buffer^84^. 16 uL of cholorform was added to each replicate and vortexed. The OD600 was measured and the permeabilized cells were diluted 1:2 in Z buffer in a 96-well plate. CPRG (2 mg/mL) was added to each well immediately before measuring the OD578 over time. OD578 was measured every 5 minutes for 50 minutes. β-galactosidase units (a.u.) were calculated by normalizing the slope of time vs OD578 measurements to the OD600 and multiplying by the dilution factor^84^.

### *Pseudomonas* Motility Assays

*P. lurida* MYb11 strains used in the swarming, swimming, and twitching assays were grown overnight in LB at 30° C. For swarming motility, 1 µL of each overnight culture was pipetted to the center of a 0.5% agar LB plate. For swimming motility, 1 uL of each overnight culture was stabbed into the agar. For twitching motility, 1 µL of each overnight culture was stabbed through 1% agar and pipetted to the bottom of the plate. The plates were incubated at room temperature for 1-2 days. The Twitching assay plates were developed using TM developer solution^85^ and surface bacteria was removed gently using a cell spreader. Images for all three motility assays were collected using a plate imaging apparatus^86^.

## Acknowledgements

The authors thank Colt Capan, Hyoungjoo Lee and Molly Soper-Hopper for sample preparation, data acquisition, and analysis of proteomics data. Proteomics experiments were conducted in the Van Andel Institute’s Mass Spectrometry Core (RRID:SCR_024903). We thank Professor Boris A. Vinatzer (Virginia Tech) for generously providing a collection of environmental *Pseudomonas* strains used in this study. Research reported in this publication was supported by the University of Michigan Natural Products Discovery Core (NPDC). The NPDC is grateful for support from the U-M Life Sciences Institute and the U-M Biosciences Initiative (RRID:SCR_023105). This work was supported by DP2DK139569.

## Author Contributions

X. Wang, K. Nauta, and N. Burton formulated the project. X. Wang, D. Gates, and M. Mechan performed the *C. elegans* screen and experiments. X. Wang and K. Nauta cloned the bacterial knock out and rescue strains. N. Chaudhary and A. Ramchandran performed the fermentation, isolation, mass spectrometry, and NMR experiments; A. Tripathi designed and supervised the natural product chemistry.. K. Sykes and C. Sakowski performed the bacterial clearance and motility assays. V. Hartwell and J. Tarango participated in screening wild type and mutant bacterial strains for their effects on dauer suppression. X. Wang and K. Nauta wrote the manuscript. All authors reviewed, edited, and approved the final manuscript.

## Supplemental Figures

**Supplemental Figure 1:**
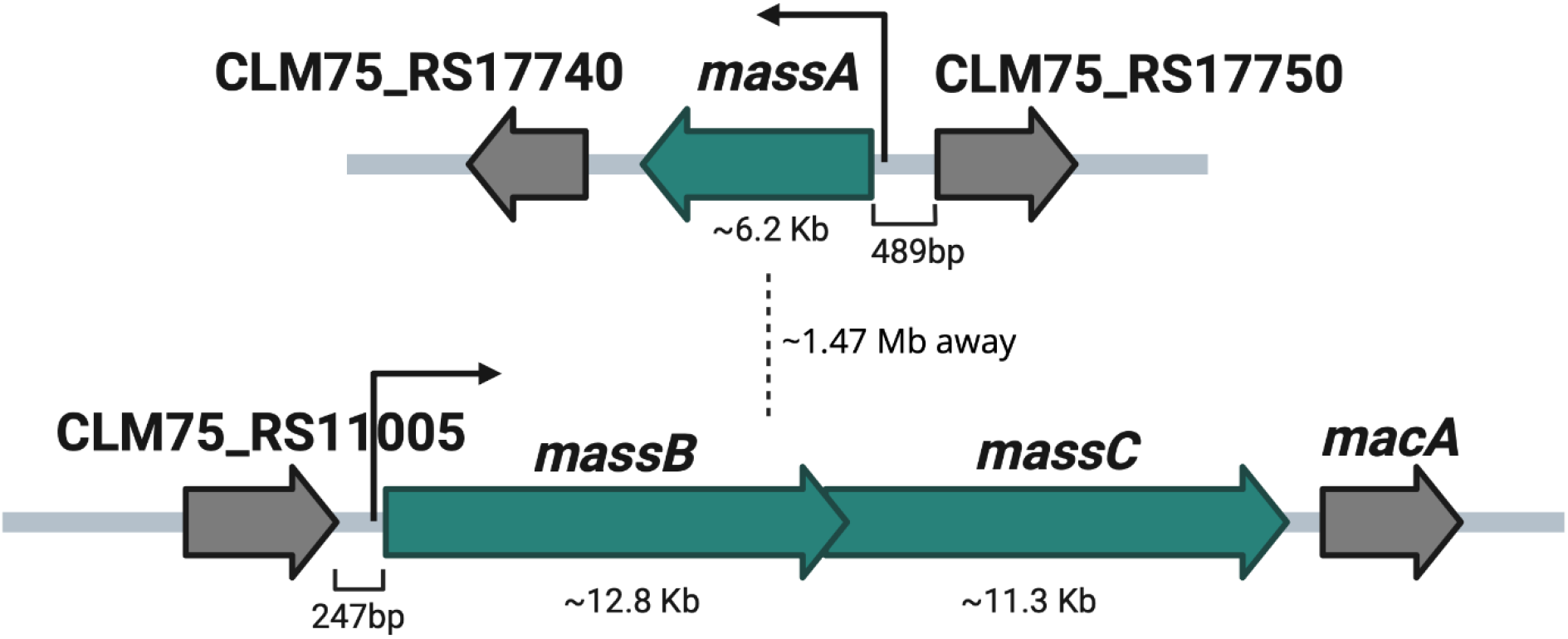
Massetolide biosynthetic gene cluster operon diagram depicting gene size and relative distance between *massA* and *massBC*.

**Supplemental Figure 2:**
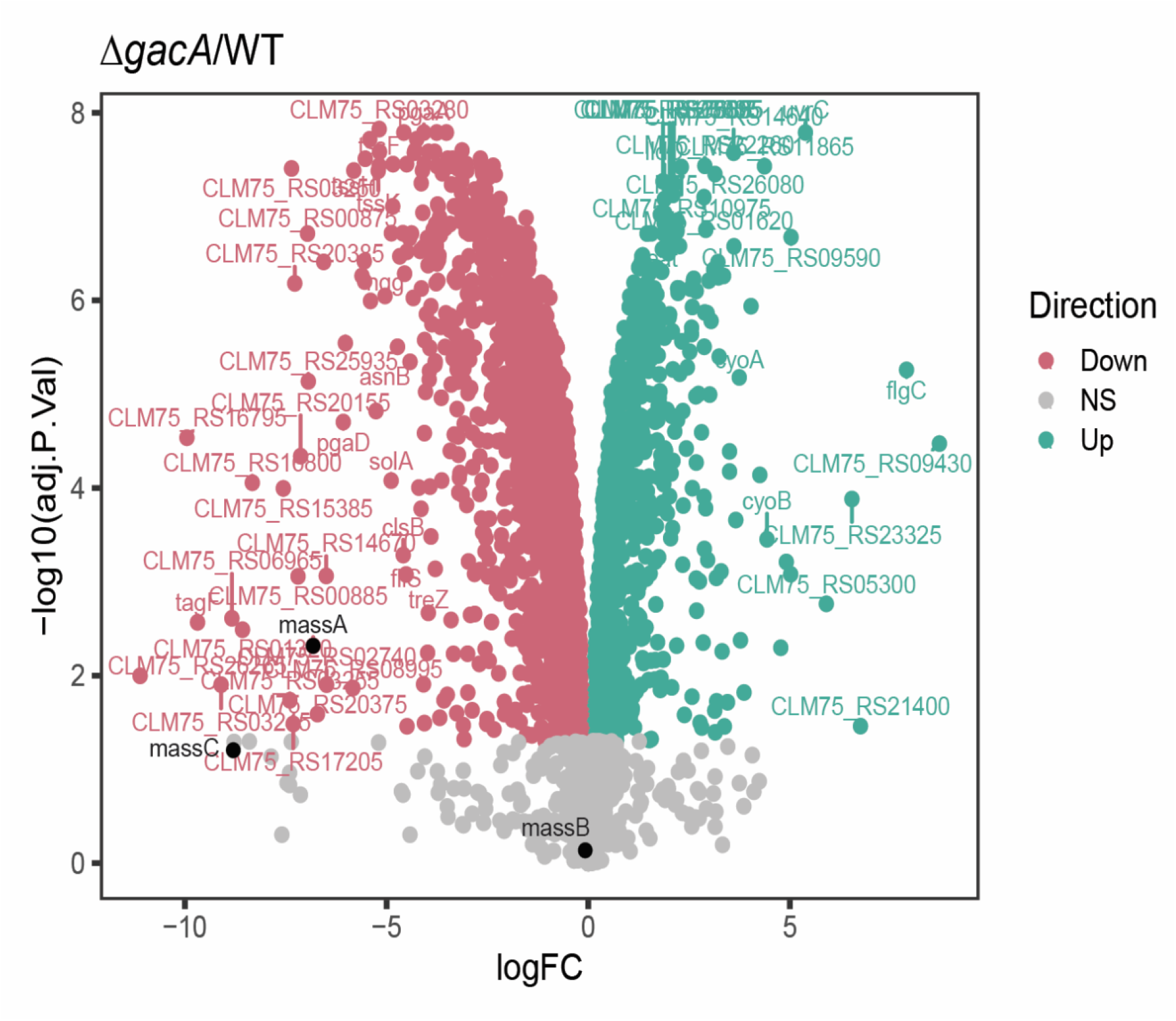
total proteomics data corresponding to figure 2E.

**Supplemental Figure 3:**
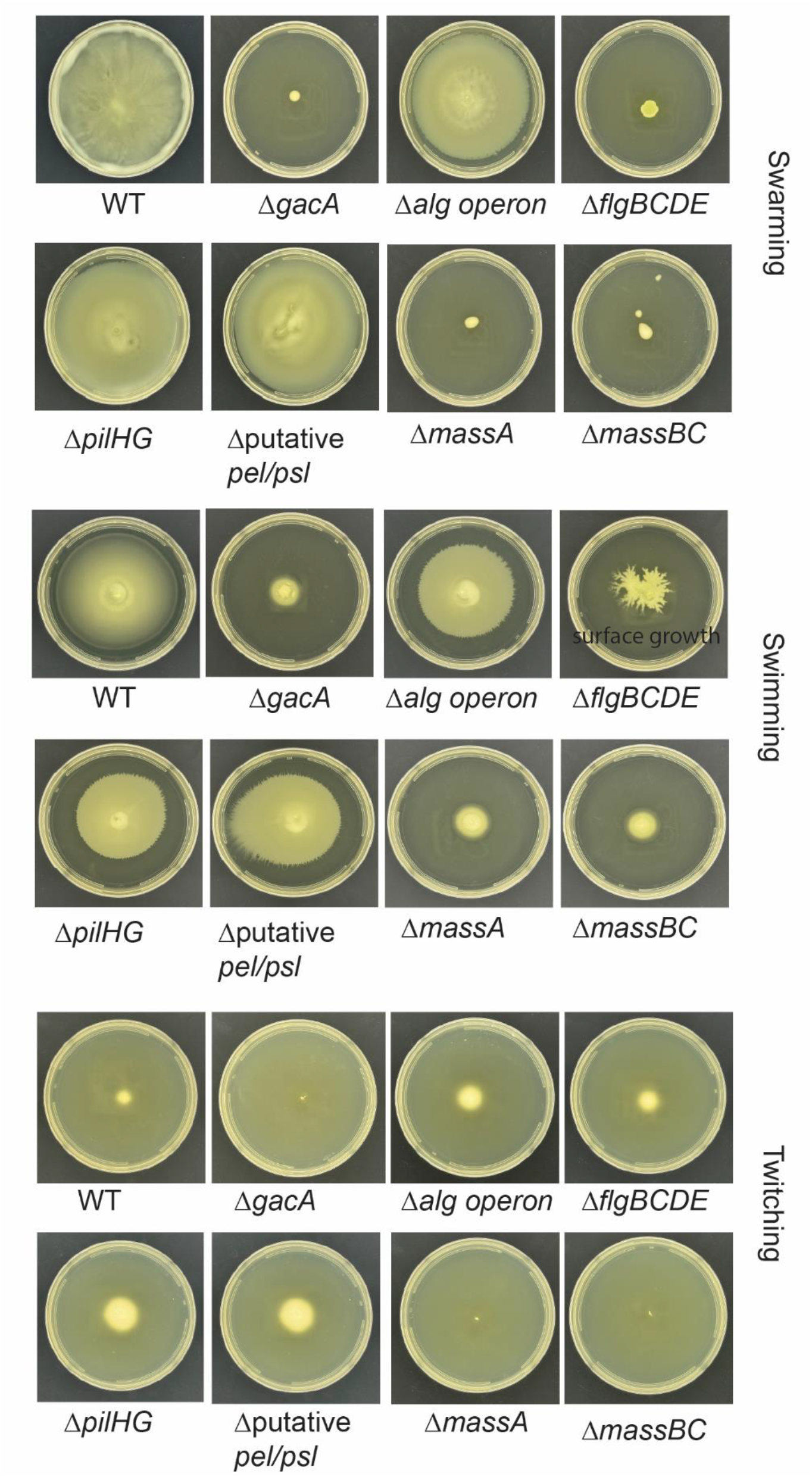
deletion of *massA* and *massBC* results in loss of swarming, swimming, and twitching motility. Pictures are representative of triplicate biological and technical replicates.

**Supplemental Figure 4:**
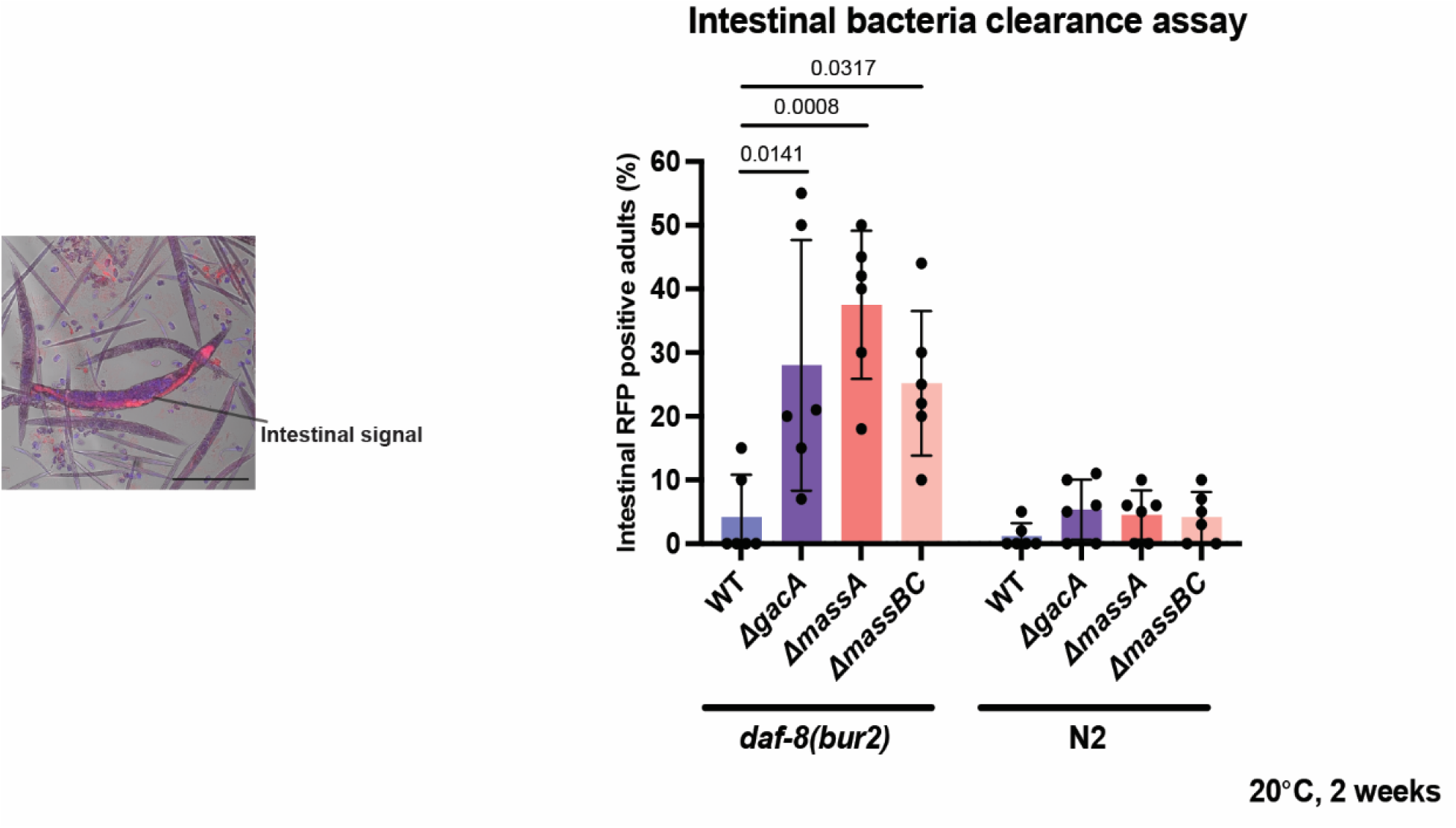
Visualization and quantification of live *P. lurida* strains in *C. elegans daf-8(bur2)* and N2 animals. Animals were grown on the indicated *P. lurida* strains at 20 °C until the young L4 stage and then transferred to NGM plates containing 50 µM FUdR and seeded with the same bacterial diet. Animals were maintained for an additional 2 weeks before collection and RNA FISH staining using the EUB338-CF610 probe targeting bacterial 16S rRNA. A representative image showing bacterial FISH signal localized within the intestinal lumen is shown on the left. For quantification, six fields of view were randomly selected from each slide, and the percentage of adult animals exhibiting detectable intestinal signal was calculated. Statistical significance was determined by ordinary one-way ANOVA followed by Sidak’s multiple comparisons testing. Error bars indicate s.d. Exact *p* values are shown on the graph; ns, not significant.

**Supplemental Figure 5:**
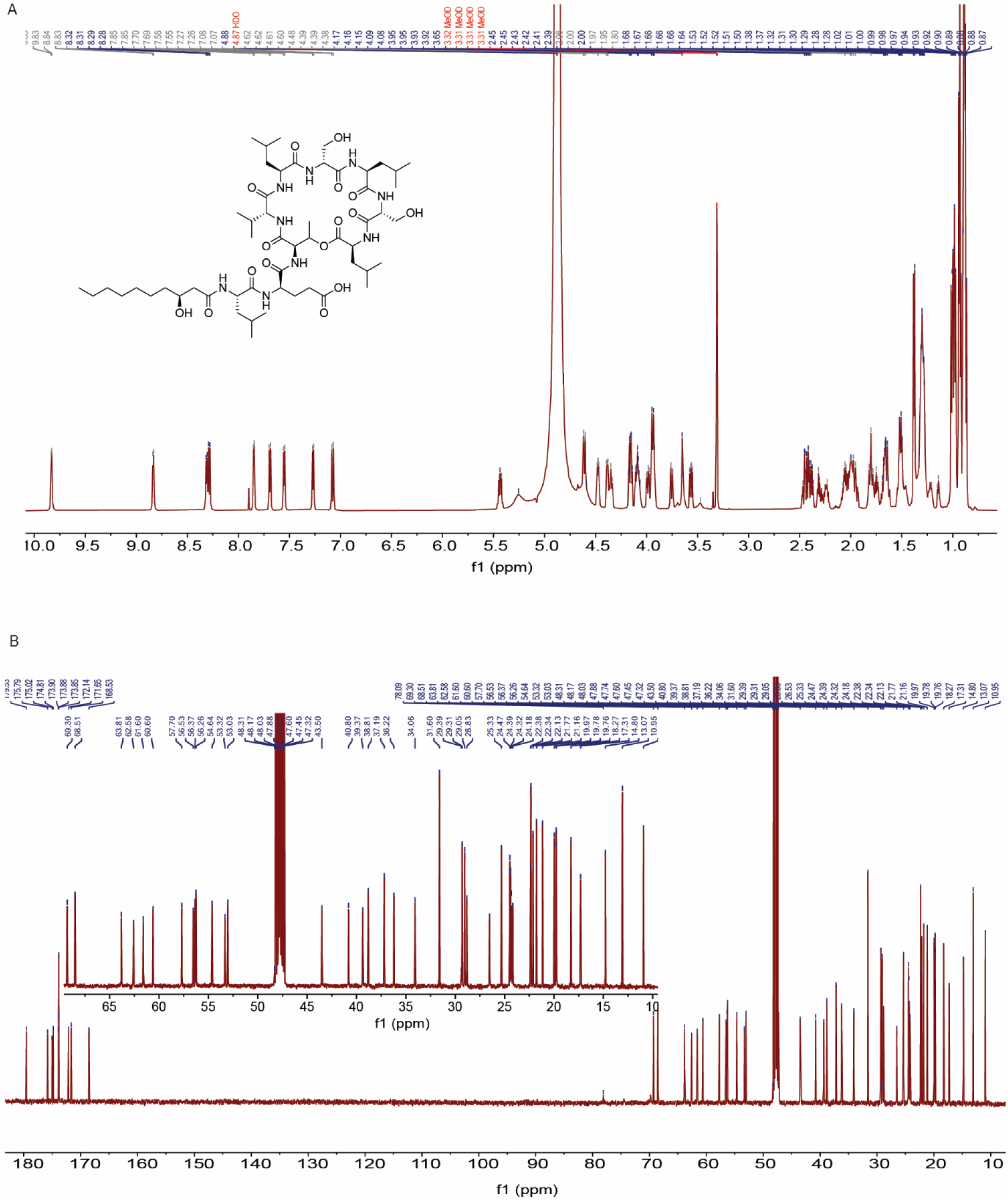
HNMR (A) and CNMR (B) corresponding to figure 3.

**Supplemental Figure 6:**
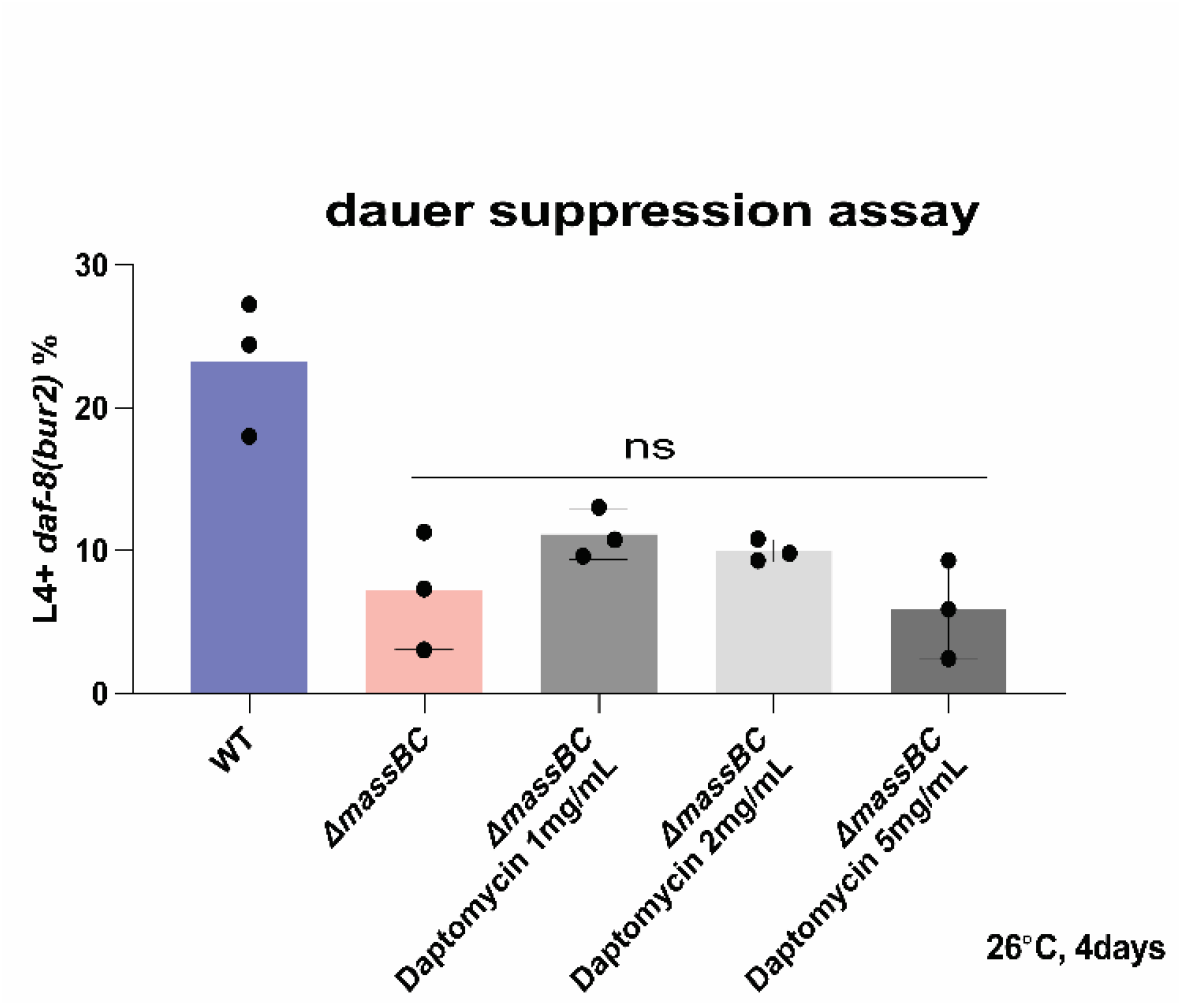
Daptomycin is not sufficient to suppress *daf-8(bur2)* dauer arrest. *daf-8(bur2)* animals were incubated on WT MYb11 and *ΔmassBC*(NOBb1018) with either vector (diH_2_O) or Daptomycin dissolved in diH_2_O (1 µg/µL, 2 µg/µL, 5 µg/µL) for 4 days at 26° C. The number of L4+ animals was counted and percentage of the total population is reported. Statistical significance was determined by ordinary one-way ANOVA followed by Sidak’s multiple comparisons testing. Error bars indicate s.d. from the mean. Error bars indicate s.d. from the mean. Exact p values are shown on the graph.

**Supplemental Figure 7:**
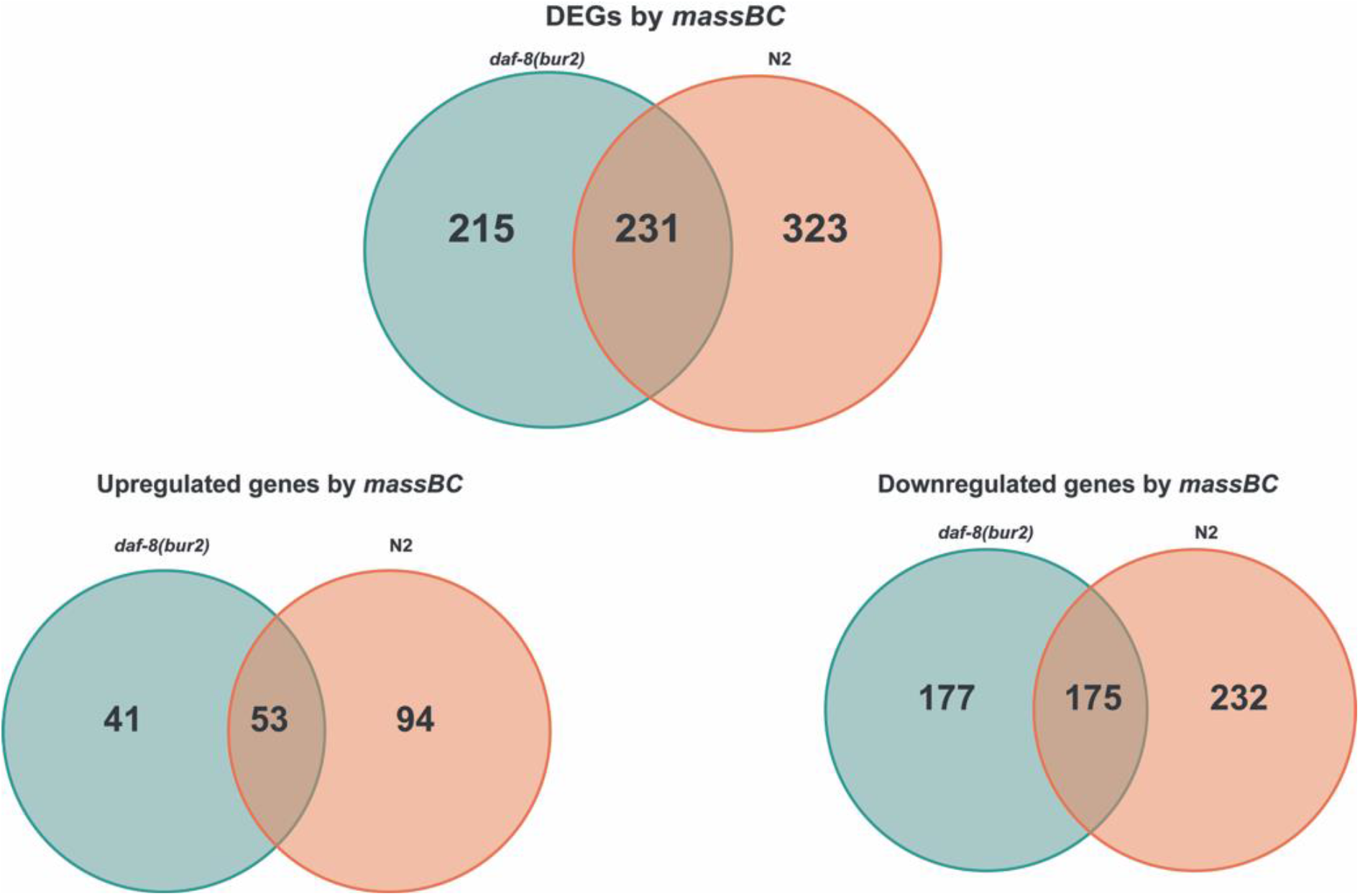
Overlapping differentially expressed genes (DEGs) in *daf-8(bur2)* and N2 animals fed Δ*massBC* versus WT *P. lurida*.

**Supplemental Figure 8:**
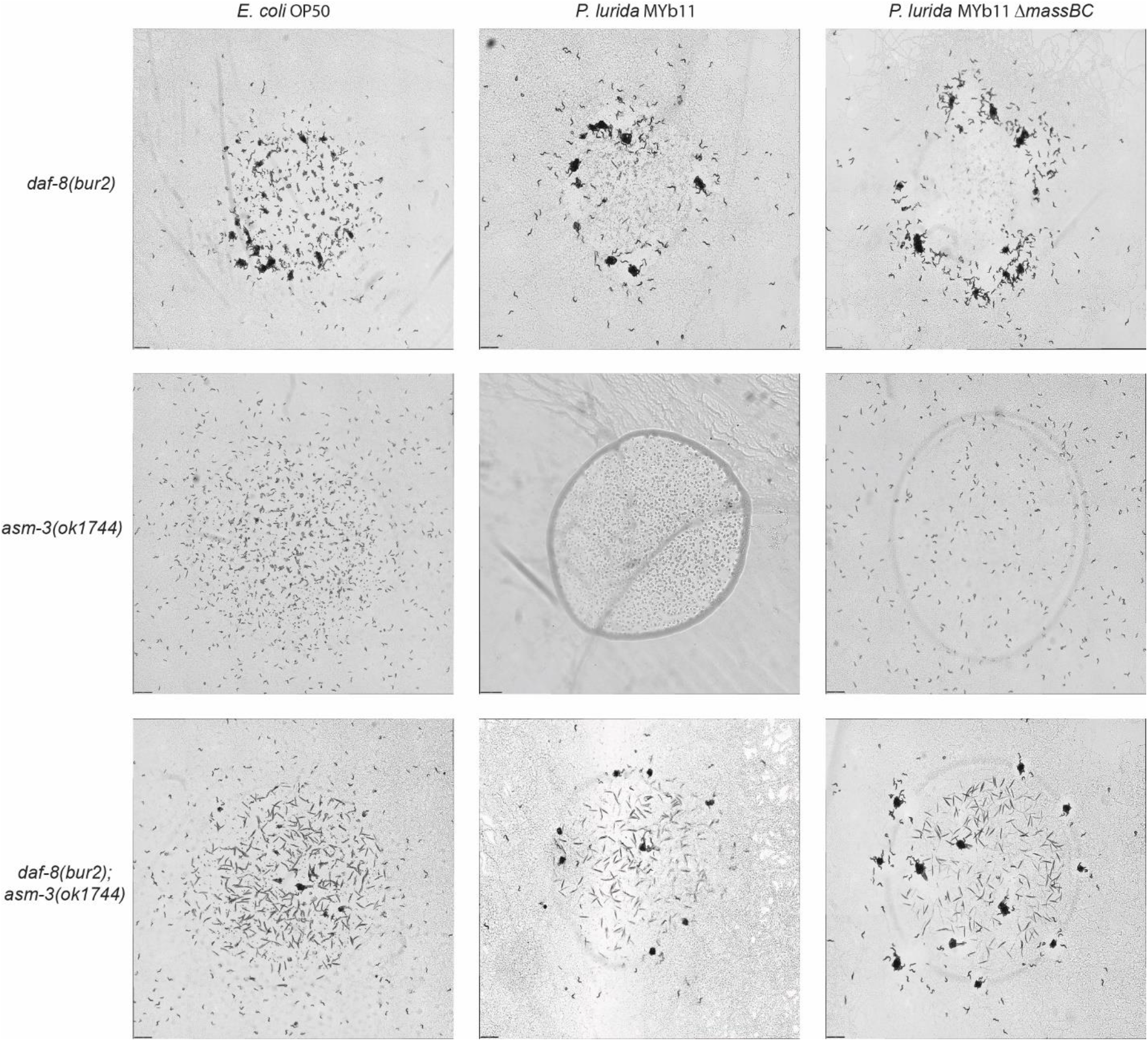
Representative images corresponding to Figure 4F of *daf-8(bur2)*, *asm-3(ok1744)*, and *daf-8(bur2)*;*asm-3(ok1744)* animals grown on *E. coli* OP50, *P. lurida* MYb11, and *P. lurida* MYb11 *ΔmassBC* (NOBb1018). Animals were incubated at 26 °C for 24 days. Experiments were performed in triplicate with approximately 500 animals per replicate.

