## Supplemental Table 4 for "Bacterial-derived cyclic lipopeptides suppress dauer arrest in *C. elegans*"

**Supplementary Table 4: List of nematode and bacterial strains used in this study.**

| **Strain** | **Description** | **Reference** |
| --- | --- | --- |
| *C. elegans* |  |  |
| N2 | *Caenorhabditis elegans* wild isolate | CGC |
| DR40 | *daf-1(m40) IV* | CGC |
| DR1572 | *daf-2(e1368) III* | CGC |
| CB1370 | *daf-2(e1370) III* | CGC |
| TJ1052 | *age-1(hx546) II* | CGC |
| JT709 | *pdk-1(sa709) X* | CGC |
| BQ1 | *akt-1(mg306) V* | CGC |
| MT1438 | *daf-1(n690) IV* | CGC |
| DR63 | *daf-4(m63) III* | CGC |
| DR72 | *daf-4(m72) III* | CGC |
| CB1393 | *daf-8(e1393) I* | CGC |
| OH7193 | *otIs181 [dat-1::mCherry + ttx-3::mCherry] III;*  *him-8(e1489) IV* | CGC |
| RB1487 | *asm-3(ok1744) IV* | CGC |
| NOB101 | *daf-8(bur2) I. T359P* | this study |
| NOB196 | *otIs181 [dat-1::mCherry + ttx-3::mCherry] III;*  *him-8(e1489) IV;*  *daf-8(bur2) I* | this study |
| NOB197 | *daf-8(bur2) I; asm-3(ok1744) IV* | this study |
| *Pseudomonas* |  |  |
| NOBb263 | *P. lurida* wild isolate from compost in Grand Rapids, MI | this study |
| MYB11 | *P. lurida* from CeMbio collection | CGC |
| NOBb335 | *P. lurida* MYB11 *ΔtssM operon (RS27440-27485)* | this study |
| NOBb441 | *P. lurida* MYB11 *ΔgacA* | this study |
| NOBb1001 | *P. lurida* MYB11 *ΔpvdM* | this study |
| NOBb1002 | *P. lurida* MYB11 *ΔphzF* | this study |
| NOBb1003 | *P. lurida* MYB11 *ΔpmbA* | this study |
| NOBb1004 | *P. lurida* MYB11 *ΔsctRST* | this study |
| NOBb1005 | *P. lurida* MYB11 *ΔRS05020* | this study |
| NOBb1006 | *P. lurida* MYB11 *ΔflgBCDE* | this study |
| NOBb1007 | *P. lurida* MYB11 *ΔflhAF* | this study |
| NOBb1008 | *P. lurida* MYB11 *ΔpilHG* | this study |
| NOBb1009 | *P. lurida* MYB11 *ΔRS09650-09655* | this study |
| NOBb1010 | *P. lurida* MYB11 *ΔR1.1 (RS00640)* | this study |
| NOBb1011 | *P. lurida* MYB11 *ΔR2.1 (RS02140-02175)* | this study |
| NOBb1012 | *P. lurida* MYB11 *ΔR3.1 (RS06675-06690)* | this study |
| NOBb1013 | *P. lurida* MYB11 *ΔR3.2 (RS06655-06670)* | this study |
| NOBb1014 | *P. lurida* MYB11 *ΔR4.1 (RS10835-10865)* | this study |
| NOBb1015 | *P. lurida* MYB11 *ΔR4.2 (RS10985-11005)* | this study |
| NOBb1016 | *P. lurida* MYB11 *ΔR4.3 (RS10960-10965)* | this study |
| NOBb1017 | *P. lurida* MYB11 *ΔR4.4 (RS10970)* | this study |
| NOBb1018 | *P. lurida* MYB11 *ΔR4.5(massBC) (RS11010-11015)* | this study |
| NOBb1019 | *P. lurida* MYB11 *ΔR5.1 (RS14700-14705)* | this study |
| NOBb1020 | *P. lurida* MYB11 *ΔR5.2 (RS14695)* | this study |
| NOBb1021 | *P. lurida* MYB11 *ΔR6.1 (RS15060)* | this study |
| NOBb1022 | *P. lurida* MYB11 *ΔR8.2 (RS17120-17130)* | this study |
| NOBb1023 | *P. lurida* MYB11 *ΔR9.1(massA) (RS17745)* | this study |
| NOBb1024 | *P. lurida* MYB11 *ΔR11.1 (RS19205)* | this study |
| NOBb1025 | *P. lurida* MYB11 *ΔR13.1 (RS27880-27895)* | this study |
| NOBb1150 | *P. lurida* MYB11 *ΔgacA P_gacA_-gacA(pNOB87)* | this study |
| NOBb1151 | *P. lurida* MYB11 *ΔmassA P_massA_-massA(pNOB88)* | this study |
| NOBb1061 | *P. lurida* MYB11 *Tn7::P_massA_'-'lacZ (pNOB12)* | this study |
| NOBb1062 | *P. lurida* MYB11 *ΔgacA Tn7::P_massA_'-'lacZ (pNOB12)* | this study |
| BAV2493 | unclassified *Pseudomonas* species | Virginia Tech University |
| BAV2591 | unclassified *Pseudomonas* species | Virginia Tech University |
| BAV2559 | unclassified *Pseudomonas* species | Virginia Tech University |
| BAV2584 | *P. antarctica* | Virginia Tech University |
| BAV2601 | *P. poae* | Virginia Tech University |
| BAV2614 | *P. trivialis* | Virginia Tech University |
| BAV2637 | *P. fluorescens* | Virginia Tech University |
| BAV2708 | *P. graminis* | Virginia Tech University |
| BAV2738 | *P. amygdali* | Virginia Tech University |
| BAV2920 | *P. tremae* | Virginia Tech University |
| BAV3042 | *P. fluorescens* | Virginia Tech University |
| BAV3129 | *P. putida* | Virginia Tech University |
| BAV3138 | *P. rhizosphaerae* | Virginia Tech University |
| BAV3141 | unclassified *Pseudomonas* species | Virginia Tech University |
| BAV3168 | *P. alcaligenes* | Virginia Tech University |
| BAV3258 | *P. putida* | Virginia Tech University |
| BAV3316 | *P. fulva* | Virginia Tech University |
| BAV3432 | *P. poae* | Virginia Tech University |
| BAV3433 | *P. cremoricolorata* | Virginia Tech University |
| BAV4009 | *P. rhodesiae* | Virginia Tech University |
| BAV4016 | *P. saponiphila* | Virginia Tech University |
| BAV4032 | *P. cedrina* | Virginia Tech University |
| BAV4044 | *P. azotoformans* | Virginia Tech University |
| BAV4094 | *P. viridiflava* | Virginia Tech University |
| BAV4119 | *P. abietaniphila* | Virginia Tech University |
| BAV4484 | *P. avellanae* | Virginia Tech University |
| CeMbio |  |  |
| BIGb0393 | unclassified *Pantoea* species | CGC |
| JUB134 | *Sphingomonas molluscorum* | CGC |
| JUB44 | unclassified *Chryseobacterium* species | CGC |
| JUB66 | *Lelliottia amnigena* | CGC |
| MYB10 | *Acinetobacter guillouiae* | CGC |
| BIGb0172 | *Comamonas piscis* | CGC |
| MYB71 | *Ochrobactrum pecoris* | CGC |
| MSPM1 | *Pseudomonas berkeleyensis* | CGC |
| BIGb0170 | unclassified *Sphingobacterium* species | CGC |
| JUB19 | *Stenotrophomonas maltophilia* | CGC |
| CEN2ENT1 | *Enterobacter xiangfangensis* | CGC |
| *E. coli* |  |  |
| OP50 | *ura-, strR, rnc-, (delta)attB::FRT-lacI-lacUV5p-T7)* | CGC |

CGC: Caenorhabditis Genetics Center

Virginia Tech University: Non-pathogenic environmental Pseudomonas from Virginia Tech

CeMbio: *C. elegans* microbiome isolates
